# Mechanistic insight into crossing over during mouse meiosis

**DOI:** 10.1101/824284

**Authors:** Shaun E. Peterson, Scott Keeney, Maria Jasin

**Author notes:** To whom correspondence should be addressed: Scott Keeney, Maria Jasin, Lead contact.

## Abstract

Characteristics of heteroduplex DNA illuminate how strands exchange during homologous recombination, but mismatch correction can obscure them. To investigate recombination mechanisms, meiotic crossover products were analyzed at two hotspots in *Msh2*^*–/–*^ mice containing homologous chromosomes derived from inbred strains. Recombination frequencies were unchanged in the mutant, implying that MSH2-dependent recombination suppression does not occur at this level of diversity. However, a substantial fraction of crossover products retained heteroduplex DNA in the absence of MSH2, and some also had multiple switches between parental markers suggestive of MSH2-independent correction. Recombinants appeared to reflect a biased orientation of crossover resolution, possibly stemming from asymmetry at DNA ends established in earlier intermediates. Many crossover products showed no evidence of heteroduplex DNA, suggesting dismantling by D-loop migration. Unlike the complexity of crossovers in yeast, these two modifications of the original double-strand break repair model may be sufficient to explain most meiotic crossing over in mice.

## INTRODUCTION

Crossing over between homologous chromosomes is essential for their proper segregation at the first meiotic division. Heteroduplex DNA, consisting of a strand from each homolog, arises during crossover formation and can encompass mismatches at sequence polymorphisms. The position and length of heteroduplex DNA have yielded insights into fungal meiotic recombination mechanisms by revealing how DNA strand information is exchanged during the recombination process. Despite decades of study, however, key questions remain about the mechanism of crossing over, particularly in mammalian meiosis where heteroduplex DNA structures of recombination intermediates remain largely untapped.

Meiotic recombination is initiated by the formation of programmed double-strand breaks (DSBs) (Keeney et al., 1997). After resection, one DNA end invades the homolog to form a displacement (D) loop, which then captures the second DNA end to form a double Holliday junction (reviewed in (Gray and Cohen, 2016)). Although they involve distinct biochemical activities, both steps—strand invasion and second end capture—result in the formation of heteroduplex DNA. Szostak and colleagues proposed that double Holliday junction resolution gives rise to two crossover and two noncrossover configurations, depending on the placement of paired nicks at each junction (Szostak et al., 1983). Subsequent studies in both yeast and mice have demonstrated, rather, that most noncrossovers arise by mechanisms distinct from double Holliday resolution (Allers and Lichten, 2001a; Cole et al., 2014; Gilbertson and Stahl, 1996; McMahill et al., 2007; Wu and Hickson, 2003). However, double Holliday junction resolution is still considered to be critical to crossover formation (Allers and Lichten, 2001b; Collins and Newlon, 1994; Schwacha and Kleckner, 1995).

In mitotic and meiotic yeast cells, mismatches in heteroduplex DNA are corrected by Msh2-dependent mismatch repair pathways. The mismatch repair machinery also suppresses recombination between non-identical sequences through heteroduplex rejection (Borts et al., 2000; Chakraborty and Alani, 2016; Spies and Fishel, 2015; Tham et al., 2016). Thus, in the absence of mismatch repair factors like *Msh2*, recombination between non-identical sequences is elevated and recombination products often show evidence of unrepaired heteroduplex DNA (Chen and Jinks-Robertson, 1999; Datta et al., 1996). In mammals, MSH2 plays similar roles in heteroduplex rejection and correction of mismatches in heteroduplex DNA in mitotic cells (de Wind et al., 1995; Elliott and Jasin, 2001), although these roles have yet to be shown in meiosis.

Genome-wide studies in budding yeast have identified thousands of crossover and noncrossover products (Anderson et al., 2015; Chen et al., 2008; Mancera et al., 2008; Mancera et al., 2011), including more recently in *msh2Δ* mutants displaying heteroduplex DNA retention (Marsolier-Kergoat et al., 2018; Martini et al., 2011). These latter studies observed complex heteroduplex DNA patterns in crossovers, leading to further modifications to the original DSB repair model including D-loop migration, double-Holliday junction migration, nick translation, template switching, and biased Holliday junction resolution.

Here, we probed mechanisms of meiotic recombination in mouse spermatocytes by defining spatial patterns of heteroduplex DNA through a comprehensive analysis of meiotic recombination products in *Msh2*^*–/–*^ F1 hybrid mice. We recovered hundreds of meiotic crossover and noncrossover products at two DSB hotspots with different levels of sequence divergence. Surprisingly, loss of MSH2 did not substantially change the recombination frequency. However, MSH2 plays an important role in mismatch correction during meiosis, allowing us to infer the presence of heteroduplex DNA. For an asymmetric hotspot in which DSB formation occurs mostly on one homolog, heteroduplex DNA was primarily observed on one side of recombinant molecules, suggesting that double Holliday junction resolution is surprisingly biased toward one of the possible configurations. Heteroduplex DNA presence was inferred in only a subset of crossover products and, moreover, these products had exchange points closer to the hotspot center than those without heteroduplex DNA. These features suggest an underlying asymmetry in the way the two DNA ends engage with the homolog and point to mechanisms to disassemble heteroduplex DNA at one of the DNA ends. We incorporate these features into a simple model of meiotic crossover formation which shares aspects with models proposed in other organisms, providing a framework for understanding fundamental meiotic recombination mechanisms from yeast to mammals.

## RESULTS

### Recombination frequencies at a symmetric hotspot are similar in wild-type and *Msh2*^*–/–*^ mice

*Msh2*^*–/–*^ and wild-type mice have equivalent numbers of MLH1 foci during meiosis (Figure S1), however, it is not clear if loss of MSH2 affects crossover placement to favor less polymorphic hotspots. To address this, we performed recombination assays at two hotspots using sperm from wild-type and *Msh2*^*–/–*^ F1 hybrid mice which contain homologous chromosomes originating from two inbred strains, A/J (A) and DBA/2J (D).The *C1* hotspot on chromosome 1 has a DSB hotspot spanning ∼200 bp, with a center of DSB activity at bp 78,590,954 in C57BL/6J (B6) mice (GRCm38; relative position = 0 bp; formerly termed the “central” hotspot; (de Boer et al., 2015; Lange et al., 2016)) (Figure 1A). The A and D mouse strains have identical sequences at the PRDM9 consensus motif to each other (and to B6) at this hotspot (Figure S2A), such that the *C1* hotspot is symmetric, with DSB formation expected to occur at equal frequency on both homologs. Within the 2.0 kb assayed, A and D have 0.6% polymorphism density, mostly from single nucleotide polymorphisms (SNPs), although a 16-bp indel is present ∼20 bp upstream of the hotspot center (Figure 1A, Figure S2A). The next closest polymorphism to the hotspot center is a SNP located 74 bp downstream.

**Figure 1.**
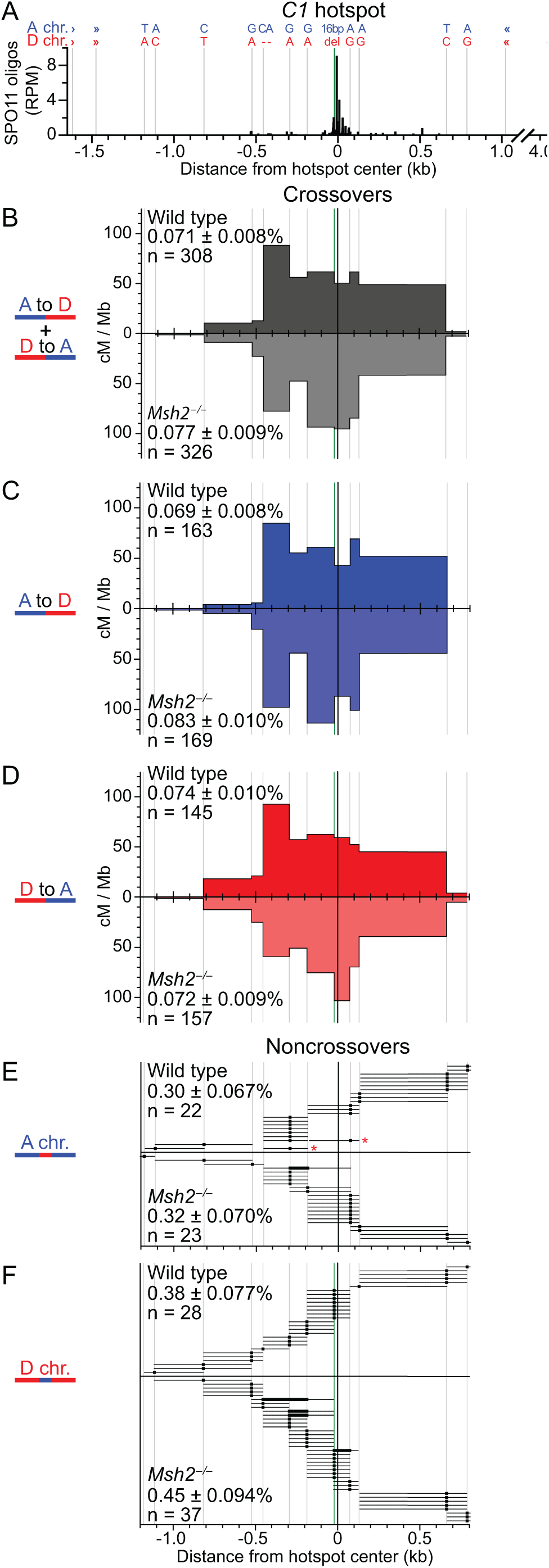
Meiotic recombination frequency and distribution are unaffected by MSH2 at *C1*—a symmetric hotspot with low polymorphism density. **(A)** The hotspot center is defined by SPO11 oligos obtained from B6 mice. Polymorphisms are indicated by vertical lines (green for the indel), with genotypes indicated above. Nested allele-specific forward and reverse primers are indicated by angle brackets. (**B–D**) Exchange interval maps for total (panel B), A-to-D (panel C), and D-to-A (panel D) crossovers. (**E and F**) Individual noncrossover gene conversion tracts, plotted on a per-well basis, for conversions on the A chromosome to D sequence **(E)**, or vice versa **(F)**. Thick dots and bars are converted polymorphisms, thin lines are the maximum possible tract lengths to the next unconverted polymorphism. Colored asterisks indicate wells showing conversion of non-contiguous polymorphisms. In B–F, Poisson-corrected frequencies are indicated ± SD; n = number of positive wells.

To specifically amplify crossover products at *C1*, nested PCR was performed on small pools of sperm DNA (∼100-200 haploid genomes per pool) using allele-specific forward and reverse primers of opposite haplotypes (i.e., A to D or D to A; Figure 1B-D, Figure S3A). To map exchange points, crossover products were probed with allele-specific oligonucleotides to identify the parent-of-origin of the polymorphisms. The 16-bp indel adjacent to the hotspot center was genotyped by PCR (Figure S2A).

We analyzed ∼9 × 10^5^ sperm genomes per mouse from two sets of littermates and recovered 308 positive wells with crossover products from wild type and 326 from *Msh2*^*–/–*^ for Poisson-adjusted frequencies of 0.071% and 0.077%, respectively (p=0.40, Fisher’s exact test; Figure 1B; Table S1). Although some variation was noted between the pairs of mice, this was unrelated to MSH2 status (Table S1). The similar frequencies suggested that neither density nor positions of polymorphisms trigger a significant MSH2-dependent suppression of recombination. As expected for reciprocal exchanges, the frequencies were similar for both the A-to-D and D-to-A orientations for wild-type and *Msh2*^*–/–*^ mice (Figure 1C,D). Crossover activity was determined by mapping the exchange interval for each crossover product (Figure 1B-D; Figure S3C). Broadly similar maps were observed for each orientation and for both orientations combined for wild-type and *Msh2*^*–/–*^ mice, as expected for a symmetric hotspot with similar DSB activity on both chromosomes.

To determine whether MSH2 affects noncrossover products at *C1*, nested PCR was performed on small pools of sperm DNA (∼15-60 haploid genomes per pool) using allele-specific forward and universal reverse primers (Figure S3B). From ∼3 × 10^4^ sperm, we recovered 50 wells with a noncrossover product from wild type and 60 from *Msh2*^*–/–*^ for Poisson-adjusted frequencies of 0.338% and 0.387%, respectively (p=0.57, Fisher’s exact test; Figure 1E,F, Table S2). More noncrossover products were identified on the D chromosome than the A chromosome, but note that the 16-bp indel was genotyped only on the D chromosome. Each conversion tract from wild-type mice contained a single polymorphism, while five tracts in *Msh2*^*– /–*^ contained contiguous conversions, which are potential co-conversions (p=0.06, Fisher’s exact test). However, two wells from wild-type mice contained two non-contiguous conversions (asterisks, Figure 1E), suggesting distinct noncrossovers, which is within the range for more than one event in the same well, as predicted assuming a Poisson distribution (Table S2).

### A highly polymorphic, asymmetric hotspot also shows similar recombination frequencies in wild-type and *Msh2*^*–/–*^ mice

Results at *C1* suggest that MSH2-dependent heteroduplex rejection either does not occur during meiosis or that the density of mismatches at *C1* is not high enough to trigger a MSH2-dependent response. To investigate the latter possibility, we examined another well-characterized hotspot, *A3* (Cole et al., 2014; Cole et al., 2010). *A3* is on the distal end of chromosome 1, with the DSB hotspot spanning ∼250 bp with the center of DSB activity at bp 160,025,733 in B6 (relative position = 0 bp; Figure 2A). Due to polymorphisms within the consensus binding site (Figure S2B), PRDM9 binds with the hierarchy D > B6 > A at *A3* (Cole et al., 2014), such that in F1 hybrids this hotspot is asymmetric, with the majority of DSBs occurring on the D chromosome. Within the 2.4 kb assayed, A and D have 1.8% polymorphism density—about 3-fold higher than the *C1* hotspot—with 41 typed polymorphisms. Other than SNPs, polymorphisms include 2-bp and 3-bp polymorphisms, and three small indels (Figure 2A, Figure S2B).

**Figure 2.**
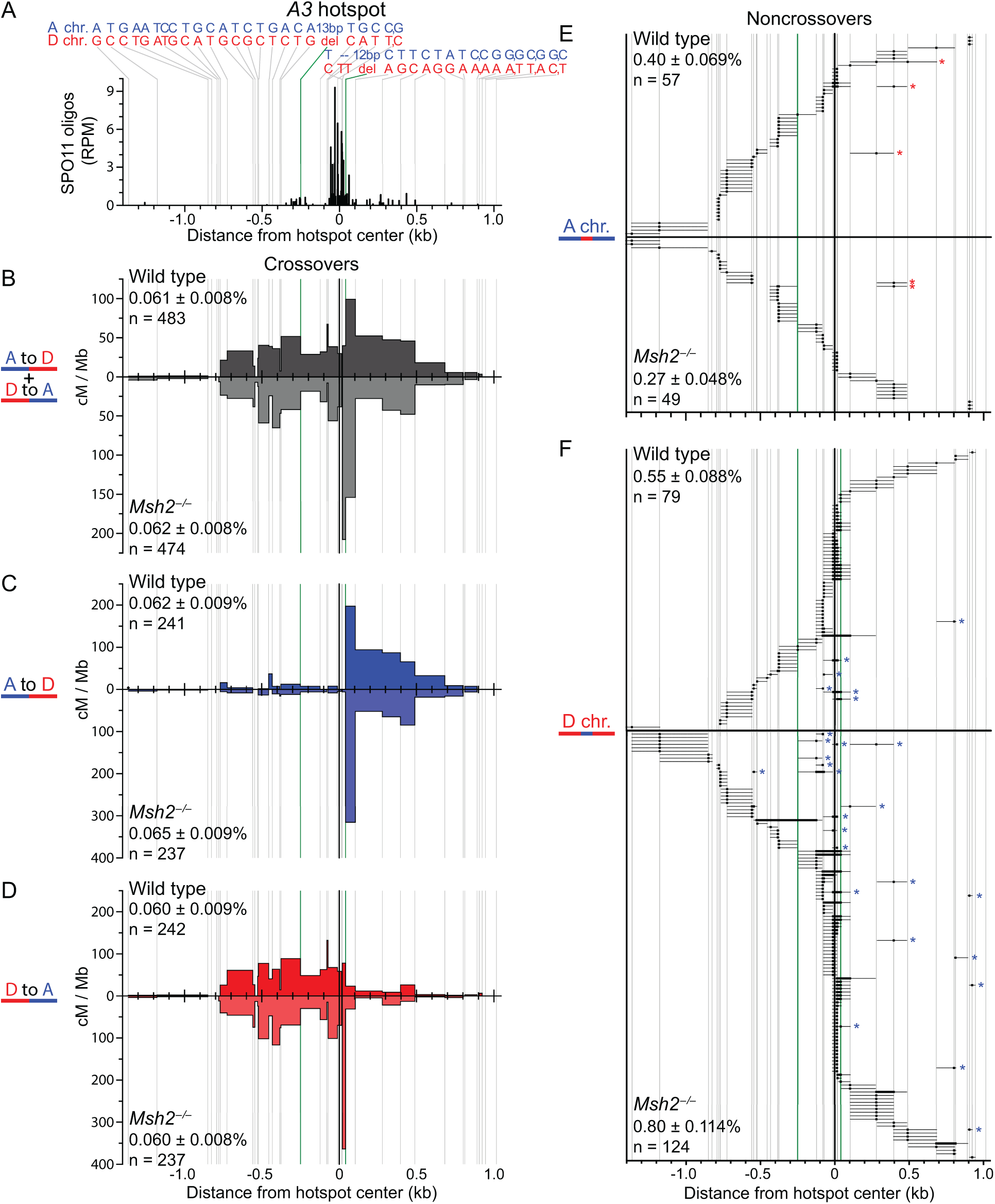
Meiotic recombination frequency and distribution are unaffected by MSH2 at *A3*—an asymmetric hotspot with high polymorphism density. **(A)** Hotspot schematic, as in Figure 1A. For genotypes shown at top, commas separate closely-spaced SNPs detected in the same allele-specific oligonucleotide. (**B–F)** Exchange point maps (B–D) and noncrossover gene conversions (E,F) are shown as described in Figure 1 legend.

We recovered 483 positive wells with crossover products from 1.7 × 10^6^ wild-type sperm genomes and 474 positive wells from 1.6 × 10^6^ *Msh2*^−/−^ sperm, among three littermate pairs, for combined Poisson-adjusted frequencies of 0.061% and 0.062%, respectively (p=0.75, Fisher’s exact test; Figure 2B, Table S1). Therefore, MSH2 also does not significantly affect crossover frequency at this highly polymorphic hotspot. A-to-D exchanges occurred with greater frequency to one side of the hotspot center (Figure 2C) and D-to-A exchanges to the other side (Figure 2D), as expected for an asymmetric hotspot with DSB formation primarily on one chromosome. Crossover maps were also generally similar between wild type and mutant, although we note more exchanges in *Msh2*^*–/–*^ in the intervals flanking the 12-bp indel at the hotspot center, possibly indicating that this indel is affecting the exchange position more strongly in the mutant.

As with crossovers, MSH2 does not significantly affect noncrossover frequency at *A3*. We identified 136 wells with a noncrossover product from 6 × 10^4^ wild-type sperm and 173 wells from 7 × 10^4^ *Msh2*^*–/–*^ sperm, among three littermate pairs, for combined Poisson-adjusted frequencies of 0.48% and 0.51%, respectively (p=0.69, Fisher’s exact test; Figure 2E,F, Table S2). For both genotypes, the noncrossover frequency was higher on the D chromosome than the A chromosome, as expected from better PRDM9 binding to the *A3* hotspot on the D chromosome. Some variability was noted for the frequencies of noncrossovers on the two chromosomes between wild type and mutant; however, variability was largely due to one of the three littermate pairs for each chromosome (Table S2).

In both genotypes, the A chromosome showed a dispersed noncrossover pattern across the interval (Figure 2E), while the D chromosome had a more clustered pattern near the hotspot center (Figure 2F), as previously observed (Cole et al., 2010). A fraction of wells contained non-contiguous conversions, suggesting two (or more) independent noncrossovers in both wild type and mutant (asterisks, Figure 2E,F), as predicted by the Poisson distribution (Table S2). Conversion of contiguous polymorphisms was also seen in noncrossovers from both wild type and *Msh2*^*–/–*^ (29% and 21%, respectively), especially on the D chromosome at the four most central polymorphisms.

### MSH2-dependent correction of heteroduplex DNA within crossover products

We identified three types of crossover products (Figure 3A; Figure S3C): simple, with one switch between parental haplotypes within a single interval (Figure 4); mixed, with both genotypes present for each polymorphism between the exchange intervals (Figure 5A, 6A); and complex, with two or more haplotype switches, possibly also including both genotypes at some polymorphisms (Figure 5B, 6B).

**Figure 3.**
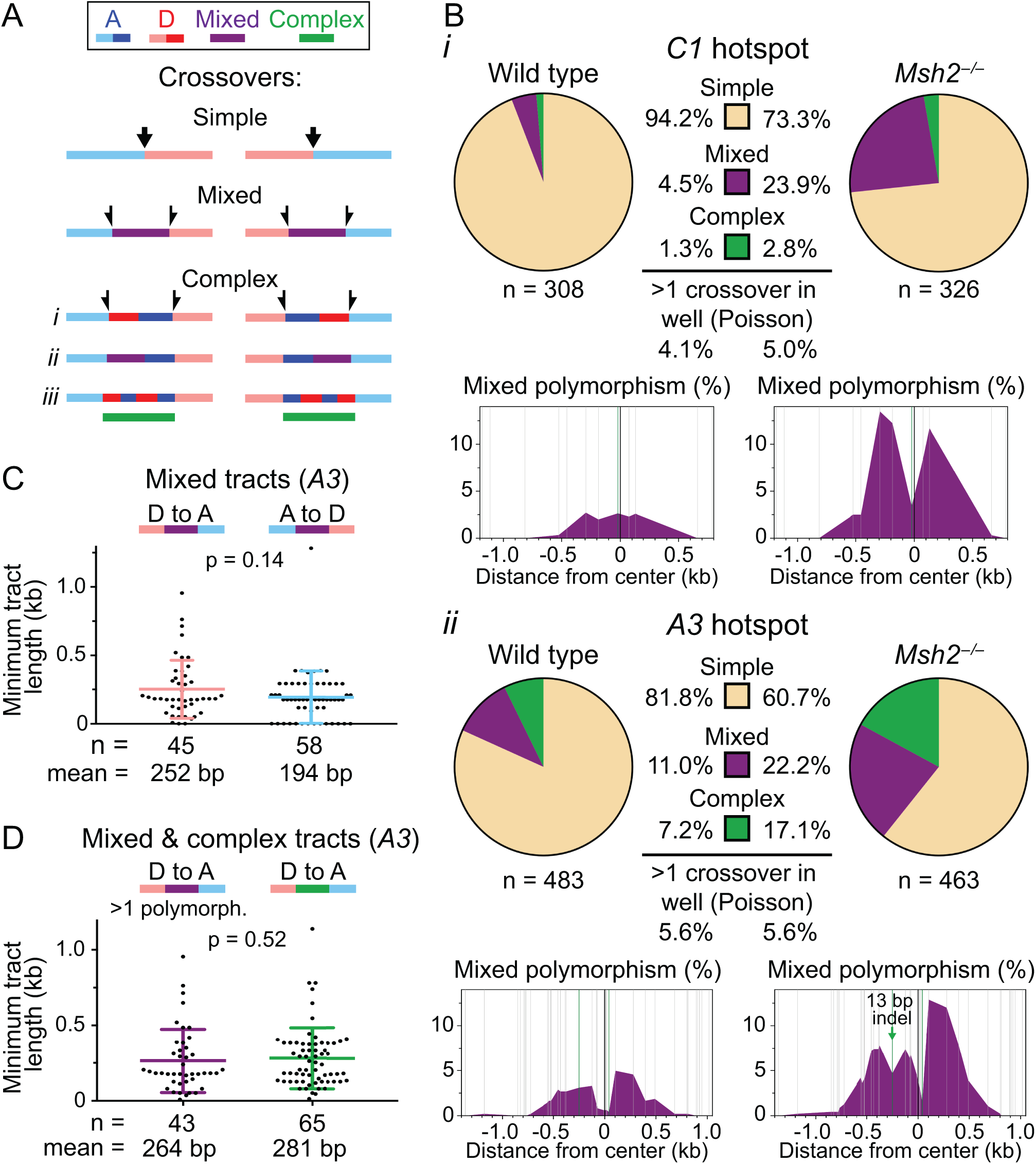
Mixed and complex crossover products are more frequent in the absence of MSH2, and likely arise from the same heteroduplex intermediate. **(A)** Crossover products were grouped into three classes: simple, mixed, and complex, with representative depictions in both A-to-D and D-to-A orientations. Exchange points, indicated by arrows, were used to calculate interval-specific recombination rates and the maps of Figures 1 and 2. Simple crossover products have a single exchange interval. For the mixed and complex crossover products, we assigned half an exchange to each interval flanking the mixed or complex tract. For complex crossover products, three sub-types were identified: (i) those containing a haplotype switch between the exchange intervals, without any mixed polymorphisms; (ii) those containing at least one mixed polymorphism; and (iii) those containing more than two haplotype switches, without any mixed polymorphisms. In this and subsequent figures, a horizontal purple bar represents a mixed tract and a horizontal green bar represents a complex tract, flanked by parental sequence (represented as light blue and pink). (**B)** The proportions of simple, mixed or complex crossover products at *C1* (i) and *A3* (ii) in wild-type and *Msh2*^*–/–*^ mice (two mice of each genotype for *C1* and three for *A3*). n = total crossover products. The Poisson-predicted percentages of positive wells expected to contain more than one crossover product are indicated below. Maps showing the fraction of mixed polymorphisms within the entire population of crossover products across the hotspot for each genotype are shown below each pie chart. **(C)** The minimum length of mixed tracts at the *A3* hotspot from *Msh2*^*–/–*^ mice was plotted separately for the D-to-A and A-to-D orientations. The lengths are not significantly different. **(D)** The minimum lengths of D-to-A mixed tracts consisting of more than 1 polymorphism were similar to the minimum lengths of all D-to-A complex tracts at *A3* from *Msh2*^*–/–*^ mice. P values in (C,D) from Mann-Whitney tests.

**Figure 4.**
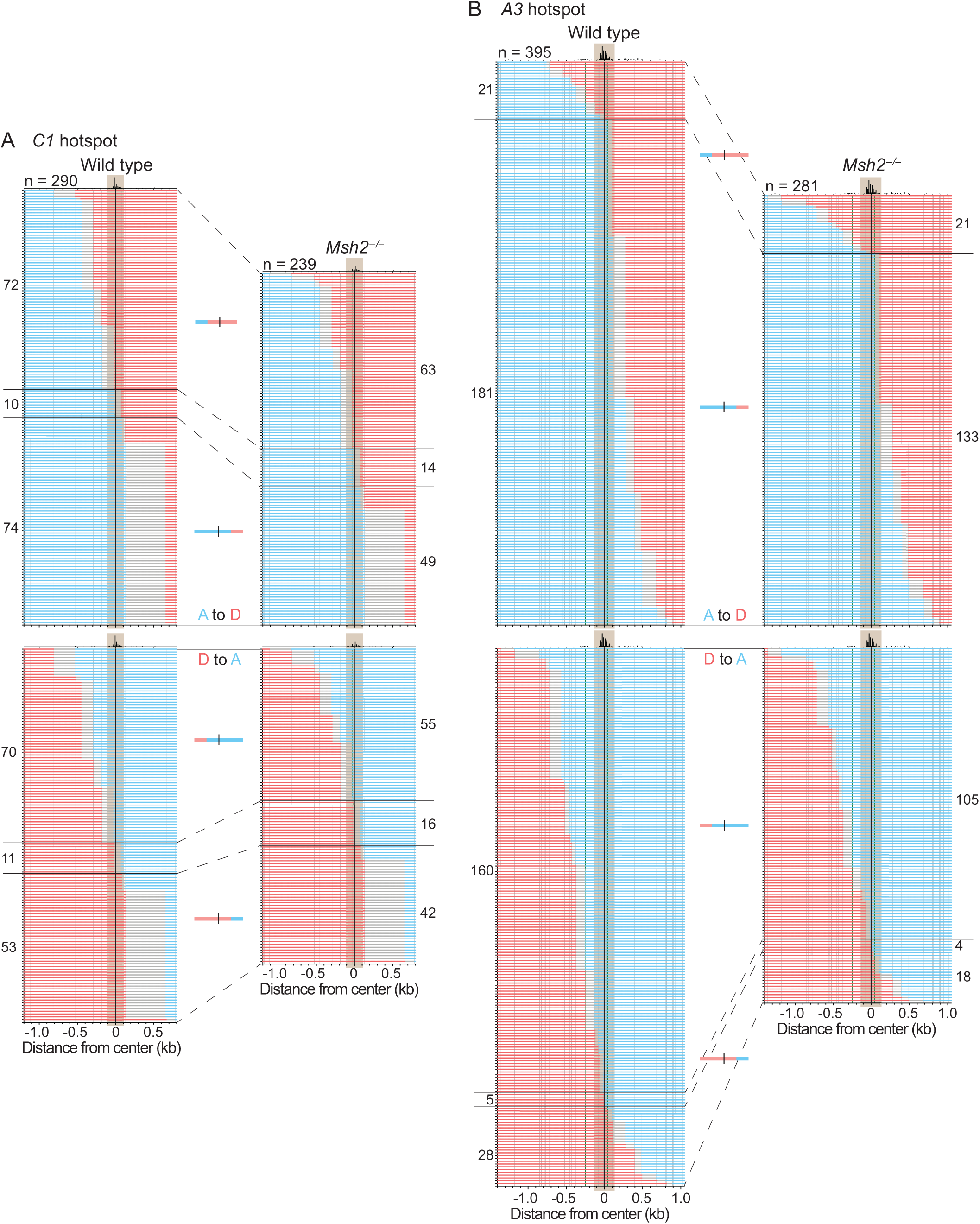
Simple crossover products. The haplotype of each crossover product at *C1* **(A)** and *A3* **(B)** is plotted horizontally, ordered by the position of exchange, with A-to-D crossovers on the top. Exchange intervals are in gray. Schematics depict four types of simple crossover products (from top to bottom): A to D with an exchange to the left of the hotspot center, A to D with an exchange to the right of the hotspot center, D to A with an exchange to the left of the hotspot center, and D to A with an exchange to the right of the hotspot center. Total numbers of simple crossovers are indicated (n). The number of crossover products in each group is indicated to the left (wild type) or right (*Msh2*^*–/–*^) of the plots. A map of SPO11 oligos (DSBs) is at the top of each stack, with the DSB hotspot shaded throughout the crossover stack (200 bp wide at *C1*, 250 bp wide at *A3*).

**Figure 5.**
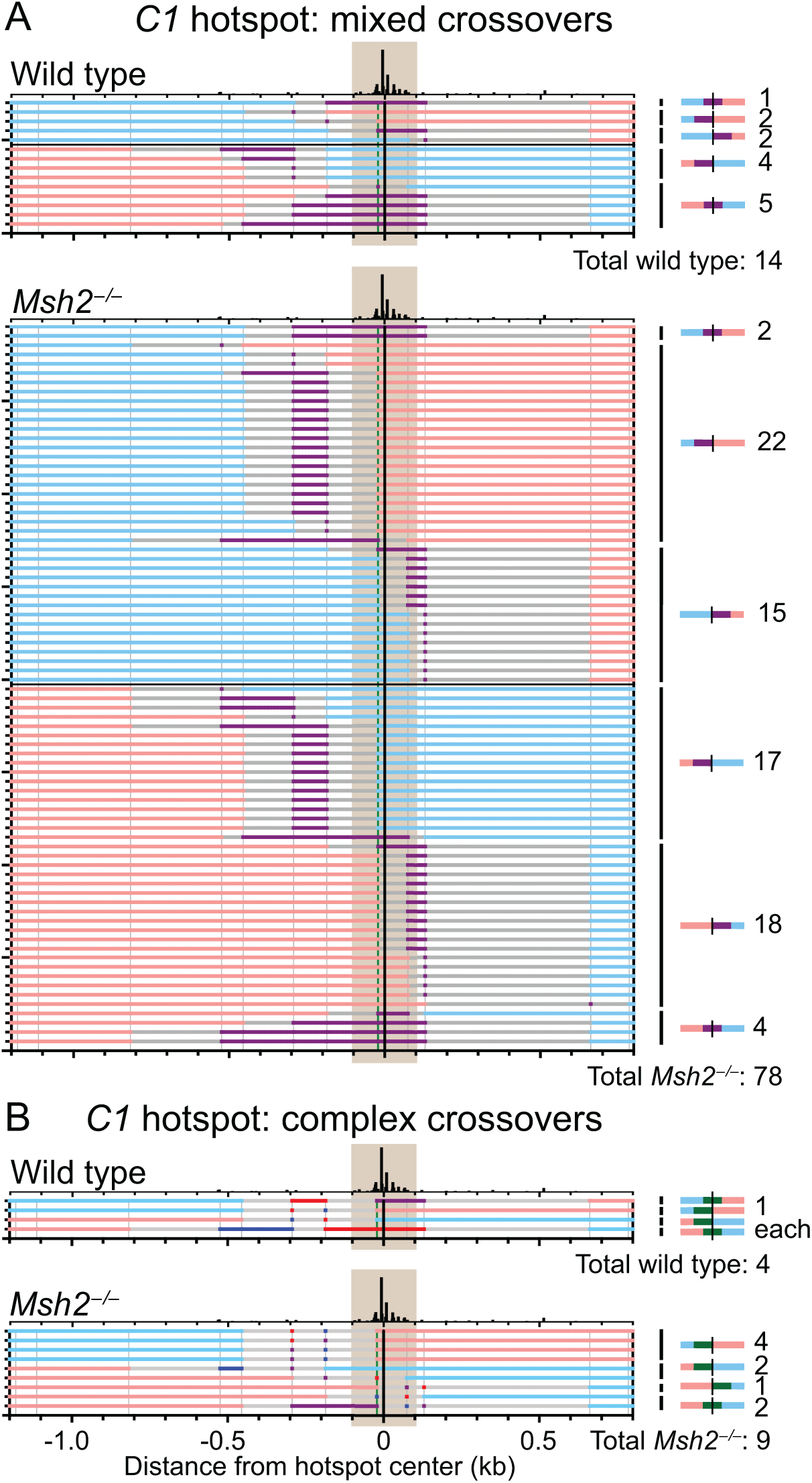
Mixed and complex crossover products from the *C1* hotspot. Wild-type and *Msh2*^*–/–*^ mixed **(A)** and complex **(B)** crossover products from *C1* are plotted as in Figure 4, with A-to-D crossover products on top. Mixed polymorphisms/tracts are indicated in purple. Schematics on the right depict the position of the mixed (purple) or complex tracts (green) within the A-to-D or D-to-A crossover products: to the left, right, or spanning the DSB hotspot, with the number of crossover products of each type indicated next to the schematic. Total numbers of mixed/complex crossover products are indicated under each panel.

Simple crossover products were the majority at *C1* in both wild type and *Msh2*^*–/–*^, but were fewer by ∼20% in the mutant (p≤0.0001, Fisher’s exact test; Figure 3Bi). This reduction primarily reflected an increase in mixed crossovers, which were 23.9% of recombinants in *Msh2*^*–/–*^. Mixed crossover products would arise from uncorrected heteroduplex DNA in a single crossover product; however, two independent simple crossovers with staggered exchange points in the same well would also appear as a mixed product (Figure S2D). For wild type, the fraction of wells expected to contain two independent recombinants assuming a Poisson distribution (4.1%) is similar to the observed percentage of wells with mixed crossover products (4.5%; p=0.69, Chi-square test). For *Msh2*^*–/–*^, two independent recombinants are expected in a similar percentage of wells (5.0%), but wells with mixed crossover products were observed at a ∼5-fold higher frequency (p≤0.0001, Chi-square test). Thus, most of the mixed crossovers in the absence of MSH2 are expected to be single crossover products with heteroduplex DNA tracts.

Broadly similar results were obtained at *A3*, although simple crossover products, while still predominating, were a smaller fraction of the total in both genotypes. The higher polymorphism density may have led to fewer simple crossover products and/or increased the ability to detect other types. Simple crossover products were again less frequent in *Msh2*^*–/–*^ (p≤0.0001, Fisher’s exact test; Figure 3Bii). In wild type, mixed crossover products (11%) exceeded the expectation for two independent crossover products in the same well (5.6%; p≤0.0001, Chi-square test), suggesting that some mixed crossover products are single crossover products with uncorrected heteroduplex DNA. In *Msh2*^*–/–*^, the fraction of mixed crossover products (22.2%) was ∼4-fold higher than the Poisson estimate (5.6%; p≤0.0001, Chi-square test), again implying that persistent heteroduplex DNA was more common in the absence of MSH2. For *A3*, the mean minimum mixed tract length from *Msh2*^*–/–*^ was 252 bp in the D-to-A orientation and 194 bp for A to D (Figure 3C, 6A).

Mixed polymorphisms were observed throughout zones extending ∼700 to 800 bp from the hotspot centers, tapering off on both sides (Figure 3B). Limits of these zones may reflect the extent of DMC1 binding (Lange et al., 2016). Mixed polymorphisms were under-represented at the hotspot centers (for example, in *A3* at the four central polymorphisms, which span ∼50 bp), and at indels, including the 13 bp indel at *A3* located 250 bp to the left of the hotspot center (Figure 3B).

Tracts with two or more mixed polymorphisms could in principle arise from heteroduplex DNA in which each strand is derived from one parental haplotype, or from heteroduplex DNA in which each strand contains switches between the two haplotypes (Figure S4A; (Hoffmann and Borts, 2005)). To distinguish these alternatives, we cloned 29 amplicons containing mixed tracts at *A3* from *Msh2*^*–/–*^ mice and determined the haplotypes in multiple subclones. Each strand within every mixed tract derived from a single haplotype (Figure S4B). Thus, switches between parental strands in heteroduplex DNA in crossover products are infrequent if they occur at all.

### Origin of complex tracts

Complex crossover products were infrequent at *C1*, but were a larger portion of products at *A3* (Figure 3B). As with mixed crossover products, *Msh2*^*–/–*^ mice had a greater proportion of complex crossover products, e.g., for *A3*, 17.1% in *Msh2*^*–/–*^ compared with 7.2% in wild type (p≤0.0001, Fisher’s exact test).

Three types of switches between parental haplotypes were observed in the complex crossover products (Figure 3A, types i–iii). For *Msh2*^*–/–*^ at *A3*, the most common (i) involved just one haplotype switch between the exchange intervals without evidence of heteroduplex retention (43 wells, 54% of complex crossover products) (Figures 3Ai, 6B). These products could not arise from two independent simple crossover products in the same well because none of the polymorphisms were mixed (Figure S3D). Where tested, these complex products yielded identical subclones, confirming the presence of a single complex crossover product (Figure S4B, complex i). Other complex crossover products (ii) had fully converted polymorphisms plus a mixed tract (30 wells, 38% of complex crossover products) (Figure 3Aii). For example in some of these, subcloning showed that one strand had multiple haplotype switches between the exchange intervals, while the other strand had no switches (Figure S4B, complex ii). The remaining six complex crossover products involved multiple switches of haplotypes without any mixed polymorphisms (8% of complex crossover products; Figure 3Aiii). Although complex crossover products were less frequent in wild type, the observed types were similar.

Complex crossover products could have originated from a continuous heteroduplex tract in which mismatched polymorphisms were corrected in an MSH2-independent manner, for example, by short patch repair (Crown et al., 2014). MSH2-independent correction appeared to be particularly robust at the 13 bp indel in *A3* (Figure 3Bii). Complex tracts were especially frequent to the left of the hotspot center in *A3* where the polymorphism density is high (Figure 6B). We compared the lengths of the mixed and complex tracts at *A3* by focusing on the numerous D-to-A crossover products from *Msh2*^*–/–*^. Because complex crossover products involve at least two switches between parental haplotypes, we only considered mixed tracts covering at least two polymorphisms. The lengths of these mixed and complex tracts were similar (Figure 3D). Thus, it is conceivable that mixed and complex tracts arise from a common heteroduplex intermediate, but with some products experiencing MSH2-independent correction. Alternatively, complex tracts could arise by other mechanisms, such as template switching, that produces tracts of a similar length.

**Figure 6.**
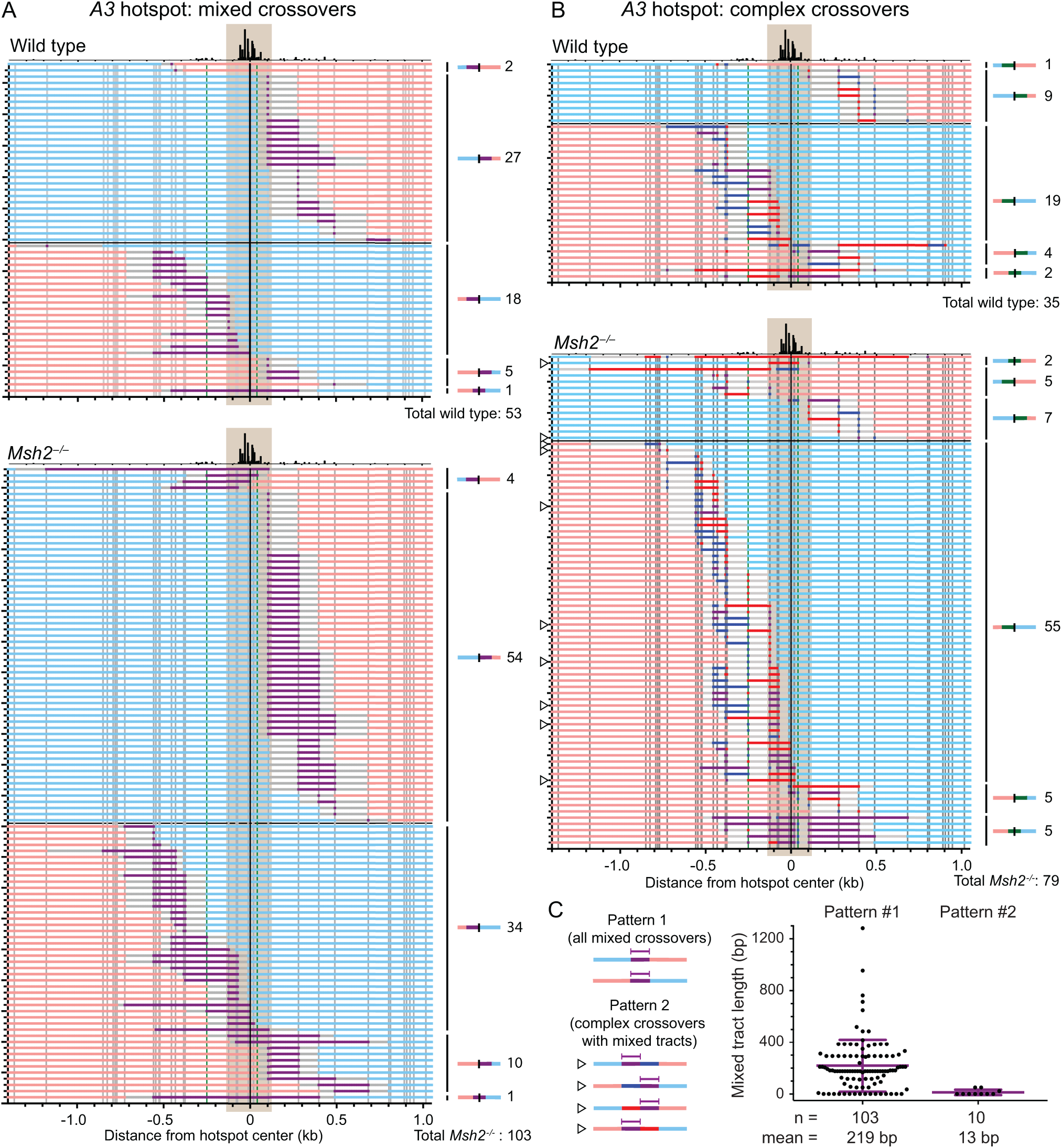
Mixed and complex crossover products from the *A3* hotspot. Wild-type and *Msh2*^*–/–*^ mixed **(A)** and complex **(B)** crossover products from *A3* are plotted as in Figure 5. Open arrowheads indicate the complex crossovers in *Msh2*^*–/–*^ that potentially conform to pattern 2 in Figure 7A. **(C)** The minimum mixed tract lengths at *A3* in *Msh2*^*-/-*^ mice are plotted for mixed crossover products which conform to pattern 1 and for complex crossover products which could potentially conform to pattern 2 (indicated by open arrowheads in B). Most mixed tracts from pattern 2 encompassed just a single polymorphism, and were on average 14.5-fold shorter than the mixed tracts from pattern 1 (p≤0.0001, Mann-Whitney test).

### Crossover patterns provide evidence for biased resolution

To gain insight into mechanisms of recombination, we grouped each type of crossover product according to the position of the exchange relative to the DSB hotspot (shaded in Figures 4-6). At the *C1* hotspot with symmetric DSB formation, a similar number of simple exchanges occurred to the left and to the right of the DSB hotspot for both the A-to-D and D-to-A orientations in wild type and *Msh2*^*–/–*^ (Figure 4A). Exchanges flanking the mixed tracts in *Msh2*^*–/–*^ also occurred on either side of the DSB hotspot, although in a few cases the mixed tract spanned the hotspot (Figure 5A). Complex crossover products were too scarce to be informative (Figure 5B).

In contrast, DSBs at *A3* form primarily on the D chromosome, leading to asymmetry in exchange positions. Simple exchanges (Figure 4B) and exchanges flanking the mixed tracts (Figure 6A) occurred primarily to the right of the DSB hotspot in the A-to-D orientation and to the left for the D-to-A orientation. The frequent complex crossover products in the D-to-A orientation showed a similar bias (Figure 6B).

In the simplest version of the canonical model for meiotic recombination, resolution of double Holliday junctions to give rise to crossovers occurs in two configurations, yielding two distinct patterns of heteroduplex DNA (Figure 7A). Considering a DSB on the D chromosome at *A3*, resolution configuration #1 (filled arrowheads) would tend to yield heteroduplex DNA to the left of the DSB hotspot in the D-to-A orientation, while resolution configuration #2 (open arrowheads) would lead to a patch of full conversion to the left of the DSB hotspot and heteroduplex DNA on the right (Figure 7A). Our finding that mixed and complex tracts in the D-to-A orientation were mostly to the left (89 of 110; Figure 6A,B) is therefore consistent with resolution primarily by configuration #1. The fewer mixed and complex tracts on the right side (15 of 110) would also be consistent with resolution configuration #1 when a DSB formed on the A chromosome (Figure S5A). The few remaining crossover products with mixed and complex tracts that span the DSB hotspot were more ambiguous but constitute only a small fraction of the total (6 of 110). As expected, an opposite bias was obtained in the A-to-D orientation: Mixed and complex tracts were primarily to the right of the DSB hotspot (61 of 72; Figure 6A,B), with few to the left (9 of 72) or spanning the DSB hotspot (2 of 72), again supporting frequent resolution by configuration #1.

**Figure 7.**
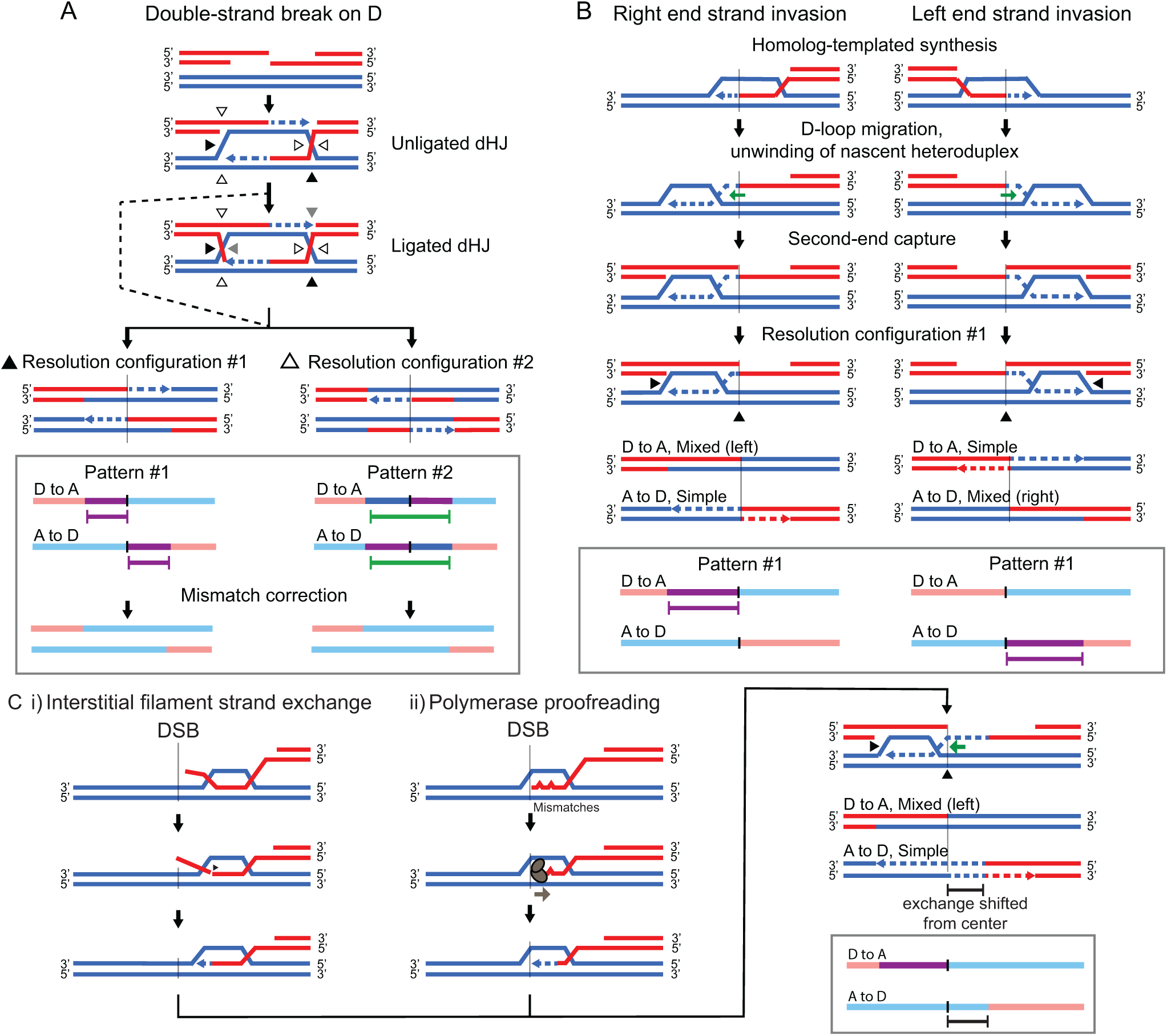
Canonical and modified models for meiotic crossover formation. **(A)** The canonical DSB repair model. A break on the D chromosome or the A chromosome (Figure S5A) leads to resection, strand invasion, and second end capture. This intermediate may mature into a fully ligated double Holliday junction (dHJ), or may be acted on by resolvases with nicks still present (dashed pathway). Single-strand cleavage at the four (ligated dHJ) or two (unligated dHJ) filled arrowheads will yield crossover pattern #1, while cleavage at the four open arrowheads will yield crossover pattern #2. The downward arrows and vertical black lines indicate the DSB position. Below these products are schematic representations of crossovers with mixed tracts indicated by purple brackets and complex tracts indicated by green brackets. Below this is the pattern that would be expected in wild type after mismatch correction. **(B)** D-loop migration model to account for abundance of simple crossovers. Initial strand invasion from each end is shown separately, yielding different crossover products. After strand exchange, repair synthesis and strand unwinding causes co-directional D-loop migration, so that the D-loop ends up on one side of the break. Subsequent second-end capture by the D-loop creates a new heteroduplex tract, and resolution of the double-Holliday junction by configuration #1 will produce one simple crossover product and one with a heteroduplex DNA tract. With frequent breaks on the D chromosome at the *A3* hotspot, this simple model can produce crossovers with the patterns we observe, namely, many simple crossovers, D-to-A crossovers with a mixed tract to the left of the hotspot, and A-to-D crossovers with a mixed tract to the right. **(C)** Alterations to the model presented in (B) that shift the simple crossover exchanges away from the hotspot center. (*i*) Strand exchange at an interstitial position of the DMC1/RAD51 nucleoprotein filament, followed by cleavage of the distal 3’ ssDNA end. Repair synthesis proceeds from the clipped end, causing the exchange in the simple crossover product to be shifted away from the hotspot center. *(ii)* The repair DNA polymerase can detect mismatches at the 3’ end of the invaded strand, which will ultimately cause the exchange to be modestly shifted away from the center in the resulting simple crossover. The crossover with heteroduplex DNA has an exchange point defined by the 3’ end of the non-invading broken strand, or, by the position/length of the D-loop prior to resolution.

In addition to considering the sidedness of the mixed and complex tracts relative to the hotspot center, we also considered the position and length of the mixed tracts within complex crossover products. Resolution configuration #2 without MSH2-independent mismatch repair is predicted to yield a patch of full conversion together with a mixed tract, which is similar in length to mixed tracts generated by resolution configuration #1. Only a few crossover products had this pattern (10 of 182; open arrowheads, Figure 6B), suggesting that this resolution configuration is rare. Moreover, mixed tracts within them typically involved a single polymorphism and thus were much smaller than tracts from mixed crossover products which encompassed multiple polymorphisms (∼14-fold smaller, Figure 6C). It is possible that MSH2-independent mismatch correction could operate on many of the products arising from resolution by configuration #2, masking their origin. However, the similar overall mean length of the mixed and complex tracts (Figure 3D) is more consistent with a common intermediate from resolution by configuration #1.

## DISCUSSION

### Similar meiotic recombination frequencies in wild-type and MSH2-deficient mice

Mammalian meiotic recombination must accommodate sequence differences between alleles without being so permissive as to risk frequent recombination between diverged, yet abundant, non-allelic repeats, which would lead to gross genome instability (Kim et al., 2016). The two hotspots we examined have 0.6% and 1.8% polymorphism density between strains, which apparently is not sufficient to trigger heteroduplex rejection during meiosis, given that both crossover and noncrossover frequencies were similar in wild-type and *Msh2*^*–/–*^ mutant mice. This degree of sequence divergence has been shown to be sufficient to substantially reduce recombination in mouse embryonic stem cells; loss of MSH2 in these cells largely restores recombination levels (de Wind et al., 1995; Elliott and Jasin, 2001; Larocque and Jasin, 2010). Thus, heteroduplex rejection may have a higher threshold for sequence divergence in meiotic cells than in mitotic cells so as not to suppress inter-homolog recombination. A greater tolerance for mismatches may be due the ability of the meiosis-specific recombinase DMC1 to accommodate mismatches within heteroduplex DNA (Lee et al., 2017; Steinfeld et al., 2019).

### MSH2 acts to correct heteroduplex DNA in mice

Our data indicate that MSH2 corrects mismatches in meiotic crossover intermediates. Mixed tracts, indicative of heteroduplex DNA retention, were obtained in the absence of MSH2 at a substantial frequency at both hotpots. Complex tracts were similar in length and position to mixed tracts, suggesting that they originated from heteroduplex DNA that was corrected in an MSH2-independent manner. At the less polymorphic *C1* hotspot, mixed and complex tracts were detected in the absence of MSH2 in 27% of crossover products, while at the highly polymorphic *A3* hotspot, they were detected in nearly 40% of crossover products. The remaining crossover products showed no evidence of a heteroduplex DNA intermediate. The detection of heteroduplex intermediates relies on amplification of both DNA strands; if only one strand were amplified a significant fraction of the time, we would underestimate the frequency of mixed tracts and, hence, heteroduplex DNA. Extrapolation to 100% amplification efficiency, however, still predicts a substantial frequency of simple crossover products (see STAR Methods – Experimental Model and Subject Details).

Another notable finding of our study was the presence of mixed and complex tracts at the *A3* hotspot in wild-type mice, above what would be expected based on Poisson correction for multiple events in a single well. Mixed tracts in wild type differ in that they more often involve only a single polymorphism (42%, 22/53) compared to the mutant (16%, 16/103; p=0.0007, Fisher’s exact test). Persistent heteroduplex DNA was also found in meiotic recombination products in wild-type yeast in a genome-wide survey (10% of all events; (Mancera et al., 2011)). In normal human sperm, complex patterns have also been detected in crossovers, albeit infrequently (Arbeithuber et al., 2015). All together, this suggests that some mismatches escape detection and repair by MSH2-dependent processes, and this may be more likely at highly polymorphic hotspots like *A3*.

Complex tracts formed in the absence of mismatch repair components have been observed in several organisms (Crown et al., 2014; Fleck et al., 1999; Guillon et al., 2005; Martini et al., 2011). *Mlh1*^*–/–*^ mice, which unlike *Msh2*^*–/–*^ mice, are defective in crossover formation as well as canonical mismatch correction, produce a small number of crossovers, and some of these contain both mixed and complex tracts, similar to what we observe in our *Msh2*^*–/–*^ mice (Guillon et al., 2005; Svetlanov et al., 2008). In fission yeast and *Drosophila* mismatch repair mutants, complex tracts are eliminated when nucleotide excision repair is also disrupted, indicating short patch repair (Crown et al., 2014; Fleck et al., 1999). Short patch repair has also been described in mammalian cell extracts (Krokan et al., 2000; Sugasawa, 2016), which could also be active in meiotic cells in the absence of MSH2. In budding yeast there is no evidence that nucleotide excision repair is linked to the formation of complex tracts in the absence of mismatch repair (Coïc et al., 2000), leading investigators to invoke alternative mechanisms, such as template switching and nick translation, to explain their origin (Marsolier-Kergoat et al., 2018).

### Crossover resolution is biased

The crossover patterns we observe suggest that double Holliday junction resolution is biased to favor one of the two hypothetical configurations (filled arrowheads, Figure 7A). In principle, this bias could arise during MLH1-MLH3-dependent resolution of either a fully ligated double Holliday junction or an unligated structure. Resolution in the former case requires four nicks, while resolution in the latter case only requires two nicks. Unlike in mouse, In which ∼90% of meiotic crossovers are MLH1-MLH3 dependent (Baker et al., 1996; Guillon et al., 2005; Lipkin et al., 2002; Svetlanov et al., 2008; Woods et al., 1999), a significant fraction of meiotic recombination intermediates in budding yeast are resolved by structure-specific nucleases (Zakharyevich et al., 2012). However, it is notable that the same resolution bias we observe in mouse has recently been demonstrated for Mlh1-Mlh3-dependent crossovers in yeast (Marsolier-Kergoat et al., 2018).

How crossover resolution bias is achieved is as yet unclear. In principle, nicks or gaps at 3’ ends in an unligated double Holliday junction would provide the asymmetry necessary to bias subsequent nicking to favor one crossover configuration (Figure 7A), as has also been proposed for budding yeast and *Drosophila* (Crown et al., 2014; Foss et al., 1999; Marsolier-Kergoat et al., 2018). It is attractive to consider that this same asymmetry could also drive resolution specifically away from generating noncrossovers, given that noncrossovers are known to occur by a distinct pathway from crossovers in mice (Cole et al., 2014). Although less parsimonious, the alternative remains possible that a crossover-to-noncrossover bias is enforced by a different mechanism than a crossover resolution configuration bias (Marsolier-Kergoat et al., 2018).

Alternatively, it has been argued that biased resolution of ligated (or unligated) substrates could be directed by proteins that promote Mlh1-Mlh3-dependent crossing over (Manhart et al., 2017). For example, the first junction formed by strand invasion and repair synthesis could accumulate proteins that stabilize this intermediate, which then direct Mlh1-Mlh3 activity in a preferred configuration, such as PCNA and Msh4-Msh5 (Manhart et al., 2017). Biochemical studies with the yeast complex suggest that it functions in a manner distinct from well-characterized structure-specific nucleases by forming higher order polymer structures (Claeys Bouuaert and Keeney, 2017; Manhart et al., 2017; Ranjha et al., 2014; Rogacheva et al., 2014), which could enforce this bias. Regardless of mechanism, directed resolution of double Holliday junctions appears to restrict the outcome of recombination to one particular crossover configuration, rather than the four possible outcomes (two crossover and two noncrossover) that were originally proposed (Szostak et al., 1983).

### Origins of simple crossover products

The original DSB repair model predicts heteroduplex DNA to be present in all intermediates giving rise to crossovers (Szostak et al., 1983) (Figures 7A, S5A). Remarkably, at the highly polymorphic *A3* hotspot, fewer than half of *Msh2*^*–/–*^ crossover products showed evidence of a heteroduplex intermediate. An MSH2-independent repair process, e.g., short patch repair, could transform a portion of heteroduplex-containing intermediates into both simple and complex crossover products (Figure S6). However, this is unlikely to explain most of the simple crossover products at *A3* since heteroduplex tracts often encompass several polymorphisms, and short patch repair appears to be inefficient as we observe many complex tracts which retain mismatched polymorphisms (Figure 6B).

Instead, our results suggest that heteroduplex DNA is retained on just one chromatid. This could be achieved by D-loop migration, which was initially proposed to explain the absence some types of aberrant segregation in yeast (Szostak et al., 1983): After strand invasion of one end, DNA repair synthesis moves or extends the D loop past the hotspot center, concomitant with disassembly of heteroduplex DNA arising from strand invasion (Figures 7B, S5B). Alternatively, strand invasion could form only very limited heteroduplex DNA. The heteroduplex DNA that gives rise to mixed tracts when MSH2 is absent would therefore primarily arise from engagement of the second end by the D loop, consistent with findings from budding yeast in which heteroduplex DNA is physically observed only in later recombination intermediates (Allers and Lichten, 2001a; Schwacha and Kleckner, 1994). Subsequent double Holliday junction resolution results in one simple crossover product and one with heteroduplex DNA. Resolution by configuration #1 is again favored, because it gives rise to heteroduplex DNA with the sidedness that we observe (Figure 7B, S5C).

In this scenario, one or more mechanisms must exist to shift exchange points at the invading end away from the hotspot center without creating heteroduplex DNA, consistent with our observations. For example, gap formation from either 3’ to 5’ end degradation (Szostak et al., 1983), flap removal after invasion from an interstitial site (Anand et al., 2014; Paques and Haber, 1997) (Figure 7Ci), and/or proofreading by the repair polymerase to degrade the invading strand at mismatches (Anand et al., 2017; Guo et al., 2017) (Figure 7Cii) would all shift the exchange away from the DSB. In these scenarios, the initial invading end and the subsequent “captured” end would undergo distinct processing steps. One prediction is that differential processing could lead to different exchange points for the simple and mixed/complex crossover products. In agreement with this, we observe that exchanges in the simple crossover products are more often outside the hotspot center compared to those from mixed and complex crossover products (Figure S7A, S7B), in that simple crossover products have exchanges which are further from the hotspot center, on average, than mixed or complex crossover products (Figure S7C).

Studies in budding yeast, including recent genome-wide studies in an *msh2Δ* mutant, have also identified crossovers in which heteroduplex DNA was present on one crossover product but not the other, often termed one-sided events (Gilbertson and Stahl, 1996; Marsolier-Kergoat et al., 2018; Martini et al., 2011; Merker et al., 2003; Porter et al., 1993). D-loop migration after strand invasion has also been invoked to explain many of these events. The complexity of meiotic products observed in yeast is enormous, leading to the proposal of a number of other variations to the canonical DSB repair model (Marsolier-Kergoat et al., 2018; Martini et al., 2011). Nonetheless, it is noteworthy that although some aspects of meiotic recombination differ substantially between yeast and mammals, mechanisms of recombination, including biased crossover resolution and D-loop migration leading to one-sided events, appear to be shared.

## ACKNOWLEDGEMENTS

The authors would like to thank Francesca Cole, Julian Lange, and members of the Jasin and Keeney labs for assistance and discussions. This work was supported by MSK Cancer Center Support Grant/Core Grant (NIH P30CA008748), a Tri-I Starr Stem Cell fellowship (SEP), R35 GM118092 (SK), and R35 GM118175 (MJ).

## AUTHOR CONTRIBUTIONS

SEP performed all of the experiments. SEP, SK, and MJ conceived of the research and designed the experiments. SEP and MJ wrote the manuscript with input from SK.

## Supplemental Information

**Figure S1.**
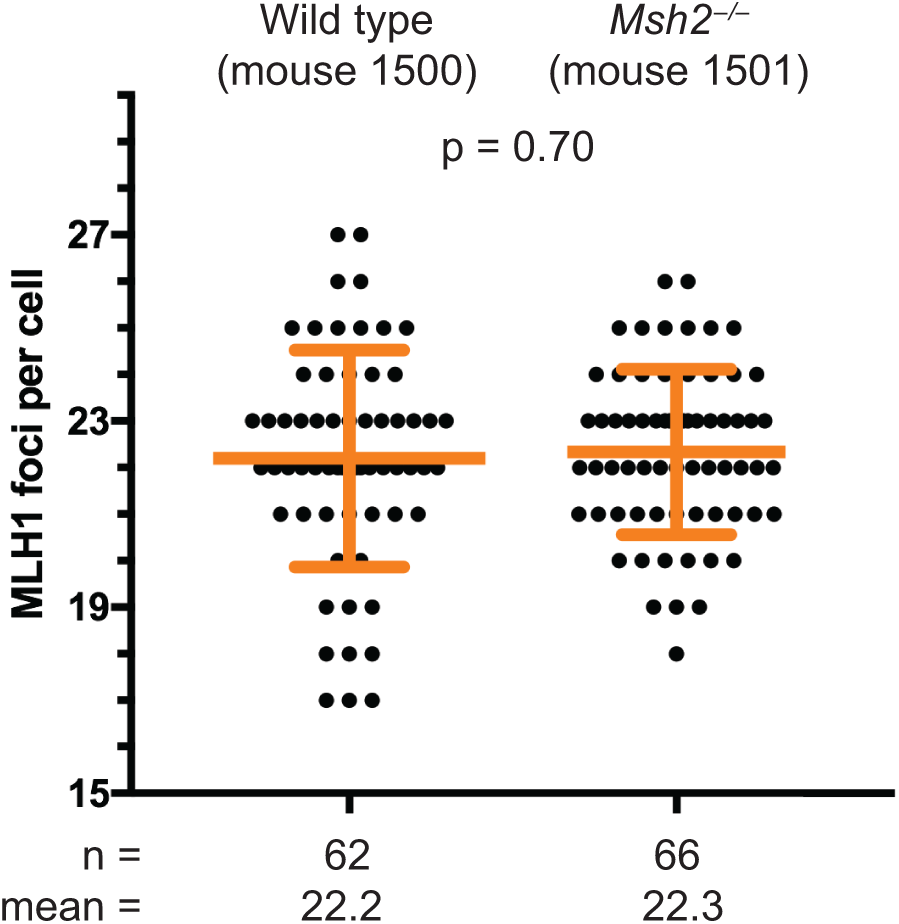
Global frequency of MLH1-positive crossovers is not affected by MSH2. Chromosome spreads from wild-type and *Msh2*^*–/–*^ F1 hybrid mice were immunostained with antibodies directed against the axial component SYCP3, and MLH1, a marker of crossover-designated sites. Each dot is the number of MLH1 foci counted in a pachytene cell. n= number of cells. P value is from unpaired t test.

**Figure S2.**
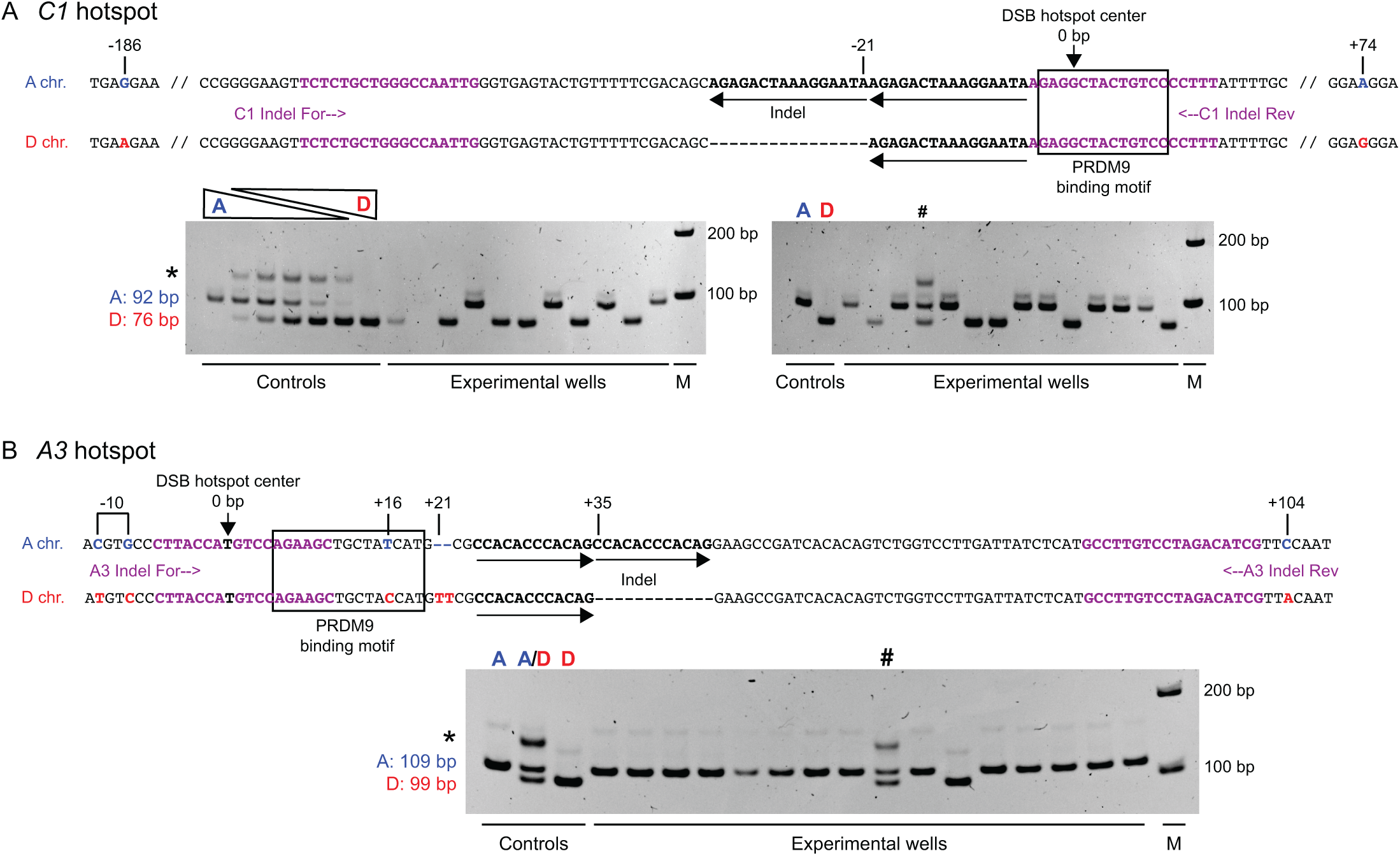
Hotspot centers, and PCR assay to genotype hotspot-proximal indels. **(A)** Partial DNA sequence of the A and D chromosomes of the *C1* hotspot, which is located in the middle region of chromosome 1. The center of DSB activity, i.e., the hotspot center, is indicated by the arrow at 0 bp. Polymorphisms are indicated by their distance from the hotspot center. The sequence matching the consensus PRDM9 binding motif is indicated with a black box (see (Yamada et al., 2017)). The genotype of both alleles of the indel cannot be determined by dot blotting, due to its repetitive nature, so a PCR-based strategy was employed on crossover products using forward and reverse universal primers (indicated in purple). Representative images from ethidium bromide-stained 5% polyacrylamide gels are shown with control reactions that are seeded with 100% A template, 100% D template or a gradient mixture of both. The sizes of the products are indicated to the left of the gel. When both alleles are present an additional band appears, indicated by the asterisk. Experimental wells contain crossover products. An experimental well which contains both alleles at the indel (a mixed crossover product) is indicated by the hash mark. For noncrossovers, PCR genotyping is not possible (see Figure S3B), however, the larger A allele at the indel can be genotyped by hybridization. **(B)** As in (A) but at the *A3* hotspot, which is located more distally on chromosome 1. The −10 oligonucleotide probe spans two SNPs, as indicated. M, DNA size marker.

**Figure S3.**
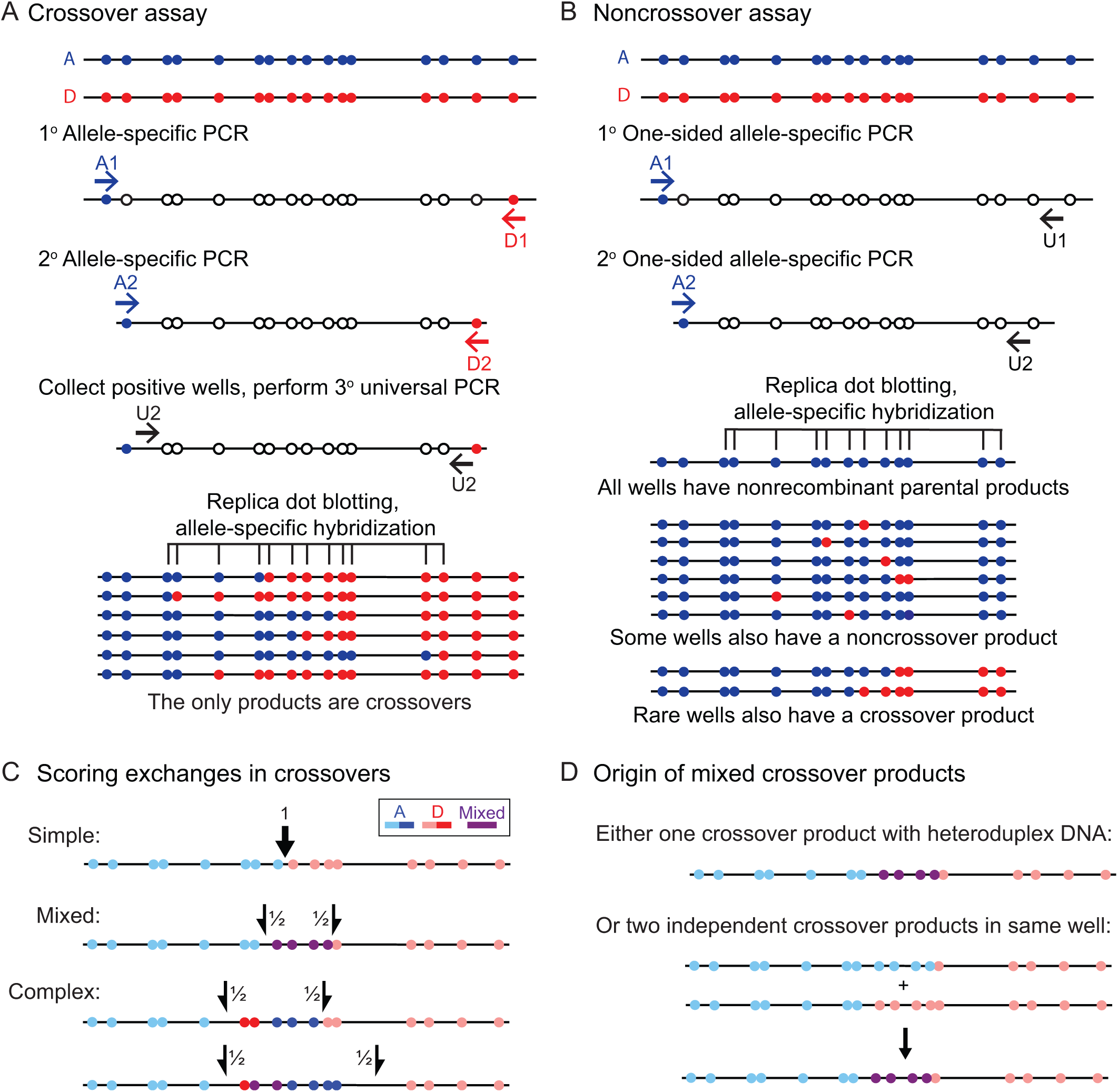
Allele-specific PCR assays to detect crossovers and noncrossovers. Sperm DNA from F1 hybrid mice from A/J (A, blue) and DBA/2J (D, red) parents are seeded in PCR reactions. (**A)** For the crossover assay (Jeffreys et al., 1998), two rounds of nested PCR are performed using allele-specific primers of opposite haplotypes (A to D is shown, D to A is not shown). In this way, the only wells giving rise to PCR products are those that contain crossovers, which are then further amplified using primers that are not allele specific (universal PCR). Internal polymorphisms are typed by replica dot blotting the products from the universal PCR onto multiple nylon membranes to be probed using allele-specific oligonucleotides (ASOs) for each polymorphism. (**B)** For the noncrossover assay (Jeffreys and May, 2004), each haplotype is amplified separately by nested PCR using allele-specific forward primers and universal reverse primers (only A to U is shown, D to U is not shown). Thus, parental molecules are amplified in each well. Those wells that also contain recombinant molecules are detected by replica dot blotting PCR products onto multiple nylon membranes and probing with ASOs. Both noncrossovers and crossovers are detected with this assay, although crossovers are not further analyzed since they are less frequent and heteroduplex DNA cannot be inferred, given the parental DNA amplification. (**C)** Simple crossover products are scored as having a single exchange interval, while mixed and complex crossover products are scored as having two exchange intervals, each given a score of ½, which flank the mixed or complex tracts, respectively. Each ½ exchange interval is bounded by a parental polymorphism (indicated by light blue and pink polymorphisms) and the first/last polymorphism in the mixed or complex tract (indicated by purple, red, or blue polymorphisms). (**D)** Mixed crossover products can derive from a single crossover product containing uncorrected heteroduplex DNA, or from two simple crossover products with staggered exchange points. The frequency of the latter case is estimated assuming a Poisson distribution.

**Figure S4.**
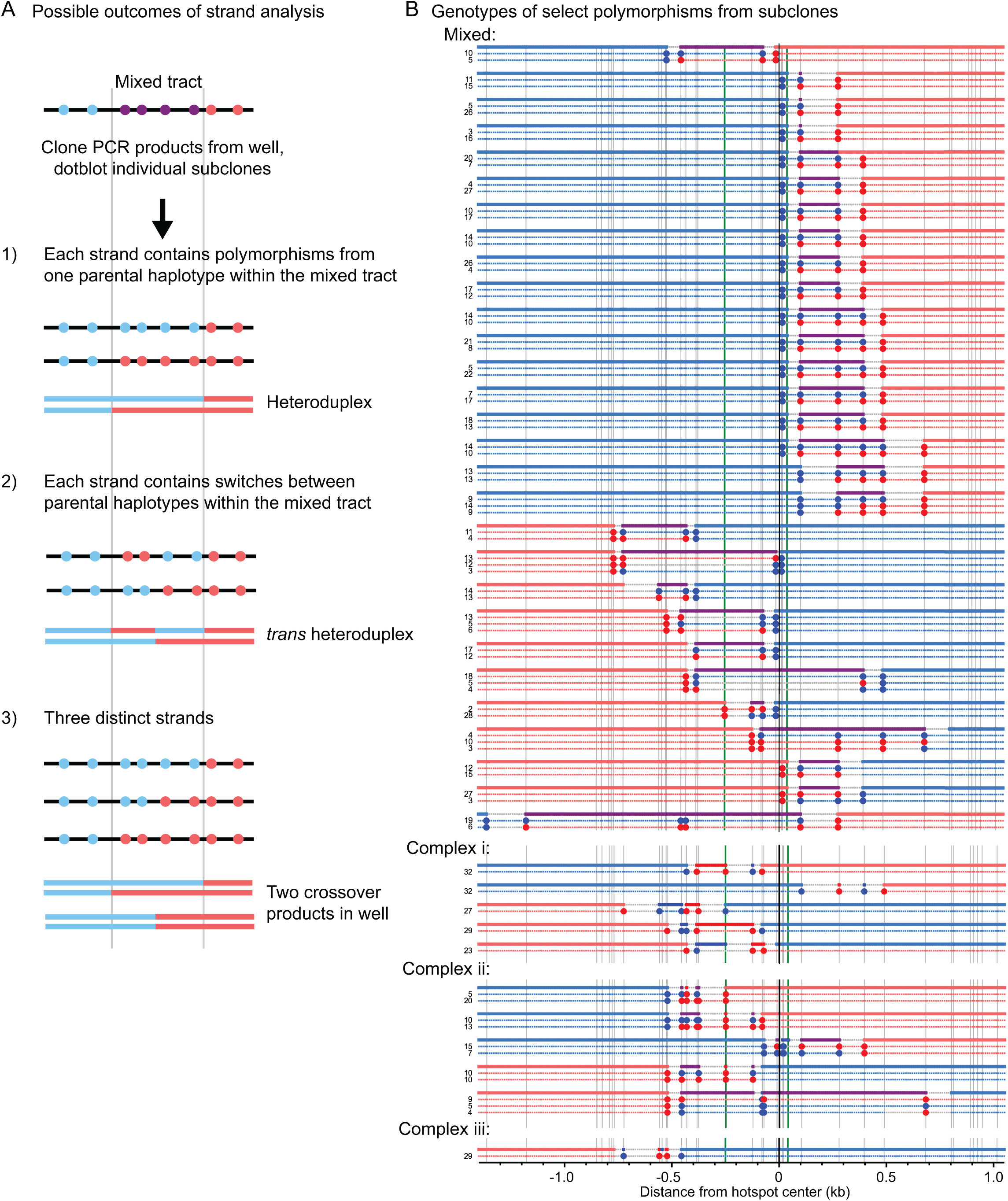
Crossover product subclones reveal a lack of *trans* heteroduplex DNA. The DNA from wells containing mixed or complex crossover products from one *Msh2*^*–/–*^ mouse was cloned, and bacterial subclones were streaked on agar plates, replica plated, lysed, and probed with the same allele-specific oligonucleotide probes as used for the crossover assay. In this way, we can distinguish the individual stands of DNA which make up a mixed or complex dot blot pattern (Cole and Jasin, 2011). **(A)** For a tract of more than two mixed polymorphisms, we can distinguish 1) simple heteroduplex DNA, 2) *trans* heteroduplex DNA, and 3) wells which contained more than two distinct strands (i.e., multiple recombinant molecules seeded in the same well). For each case, only the most likely scenario is indicated but other possibilities exist. Two crossover products with simple exchanges in the same well will appear to have simple heteroduplex DNA (1), the frequency of which is estimated assuming a Poisson distribution. **(B)** Haplotype phasing information for recombinants with mixed or complex tracts. The original dot blot hybridization pattern for each well is indicated with thick colored lines. Polymorphisms typed in the subclones are represented as blue or red filled circles. Only a few informative polymorphisms were typed for each clone. The genotype for the rest of the strand is inferred and represented by the thin dotted blue and red lines. Gray lines represent the inferred interval of exchange between parental and the mixed (or complex) tract. To the left are the numbers of individual subclones recovered with the indicated pattern. Patterns recovered from only one subclone are not shown. Mixed crossover products were never found to have strands which switched parental haplotypes within the mixed tract (class 2 in panel A), indicating a lack of *trans* heteroduplex within crossovers. Complex crossover products without mixed polymorphisms (types i and iii in Figure 3A) all yielded only identical subclones, confirming that complex crossover patterns are not the result of two independent crossover products seeded in the same well. Complex crossover products with mixed polymorphisms (type ii in Figure 3A) yielded two distinct subclone patterns. Type ii may arise from correction of one or more mismatched polymorphisms within a heteroduplex tract, whereas types i and iii may arise from correction of all mismatched polymorphisms.

**Figure S5.**
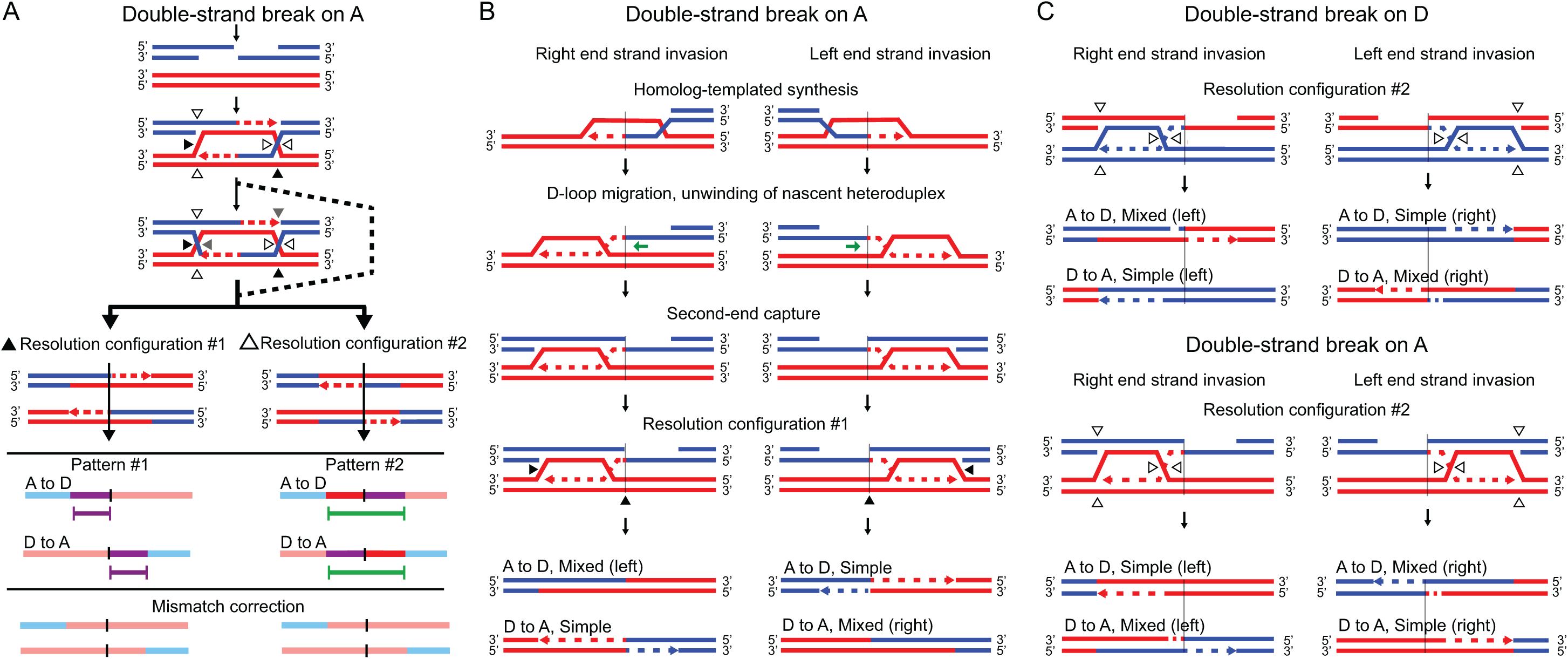
Other crossover models. **(A)** Companion to Figure 7A, but with an initiating DSB on the A chromosome. **(B)** Companion to Figure 7B, but with an initiating DSB on the A chromosome. **(C)** Companion to Figure 7B and Figure S5B, but with resolution configuration #2 rather than resolution configuration #1. Note that with D-loop migration, resolution configuration #2 does not produce complex crossovers as in Figure 7A. However, the sidedness of crossovers obtained with D-loop migration and resolution configuration #2 is opposite of what we observe. For example, with the majority of DSBs on the D chromosome, D-loop migration and resolution configuration #2 would produce A-to-D crossovers with the mixed tract on the left, and D-to-A crossovers with the mixed tract on the right. These events are under-represented in our dataset.

**Figure S6.**
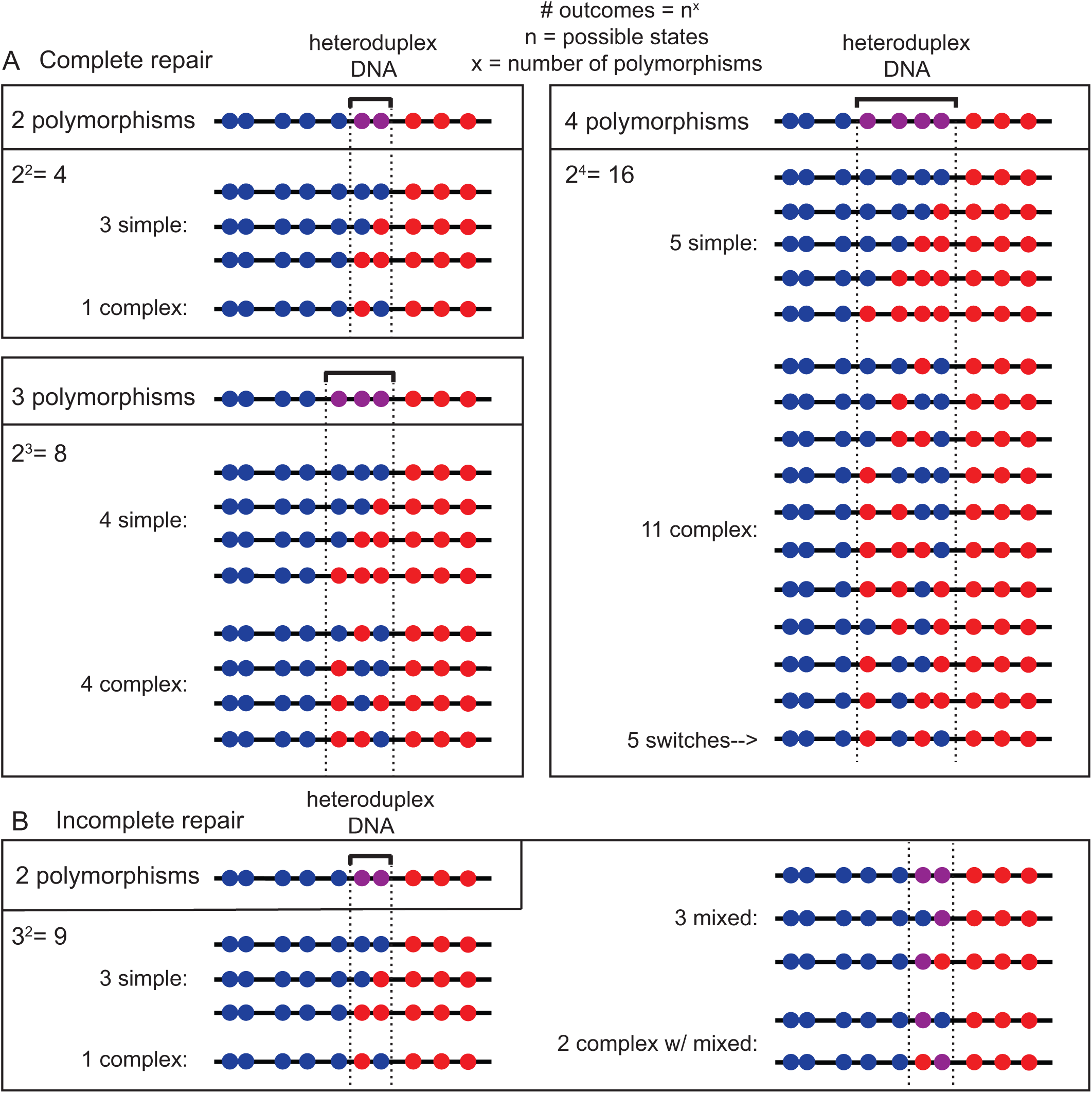
The majority of simple crossovers at *A3* are unlikely to arise from MSH2-independent repair of mismatches. In principle, a proportion of heteroduplex-containing intermediates can be transformed into either simple or complex crossovers via MSH2-independent repair. For a heteroduplex tract with *x* number of polymorphisms, the number of possible states, *n*, is 2 (for both possible genotypes) if repair is assumed to be complete. If repair is not complete, then *n =* 3 (genotype 1, genotype 2, or unrepaired). The number of possible outcomes, and types of outcomes, can be predicted, assuming that each polymorphism is repaired (or not) randomly and independently of its neighbor. **(A)** If each mismatch is corrected independently in either direction, then the fraction of simple repair products is inversely proportional to the number of polymorphisms within the heteroduplex tract. In the case of a heteroduplex tract containing two polymorphisms, 75% of repair products are expected to be simple and only 25% complex, while for three polymorphisms, 50% are expected to be simple and 50% complex. With four or more polymorphisms included in the heteroduplex tract, the majority of repair products are complex. **(B)** The estimates in (A) include only two possible outcomes – mismatch repair at each polymorphism without retention of heteroduplex DNA; thus, it overestimates the occurrence of simple events. For example, if a mismatch is equally likely to be repaired in either direction or not repaired at all, a crossover with two polymorphisms in the heteroduplex tract would be expected to result in a simple event only 33% of the time, rather than 75% of the time if repair was complete. At *C1*, mixed tracts in the *Msh2*^*–/–*^ mutant encompass on average 2 polymorphisms (2.0 for A to D and 2.4 for D to A) (Figure 5A). Given that mixed tracts comprise a much greater fraction of events (23.9%) than complex events (2.8%) (Figure 3B) and most of the complex events retain mixed polymorphisms (Figure 5B), MSH2-independent correction of mismatches is unlikely to be highly robust. At *A3*, mixed tracts encompass an average of 2.8 polymorphisms in the A-to-D orientation and 6.1 polymorphisms in the D-to-A orientation (Figure 5A). Thus, especially in the latter case, only a fraction of simple events are expected to arise from MSH2-independent correction if all polymorphisms are treated independently. However, some closely-spaced polymorphisms may be co-repaired, effectively lowering the number of repair outcomes. In particular, considering closely-spaced polymorphisms within the context of a complex tract which has additional flanking complex polymorphisms, repair of each frequently occurs in the same direction, rather than the expected 50% if they are independently repaired: i.e., for D to A, −455/-432, 5 of 6; −384/-376, 10 of 12; −80/-72, 4 of 4.

**Figure S7.**
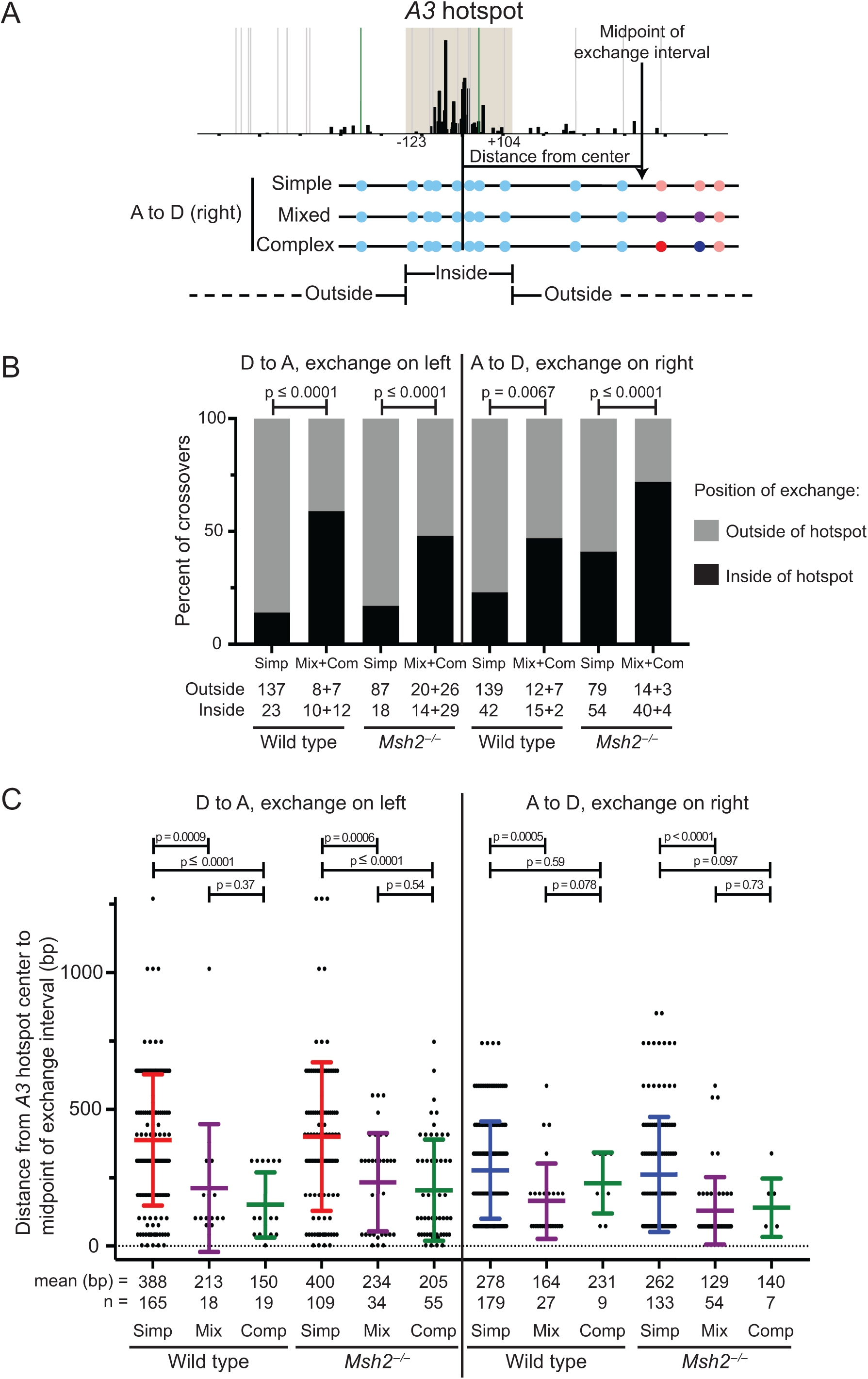
Exchanges of simple crossovers are further from the hotspot center than those containing heteroduplex DNA. **(A)** We define the DSB hotspot based on SPO11 oligo maps obtained from B6 mice. The *A3* hotspot is 250 bp wide, which places the left boundary at the −123 polymorphism and the right boundary at the +104 polymorphism. Crossover exchanges *in Msh2*^*–/–*^ mice at *A3* were categorized as falling inside or outside of these boundaries, separately for simple, mixed and complex crossovers. To plot the distance of the exchange from the center of the hotspot, we used the midpoint of the exchange interval as the position of exchange. **(B)** The percentage of simple, or mixed and complex (grouped together) crossovers with exchanges that fall inside or outside of the *A3* hotspot are plotted, separated by orientation. The number of crossovers is shown below the graph. The difference between the simple crossovers and the heteroduplex DNA-containing crossovers is significant in wild-type and *Msh2*^*–/–*^ mice for both A-to-D and D-to-A orientations (Fisher’s exact test). **(C)** The distance from the *A3* hotspot center of each crossover exchange (as defined in (A)) was plotted for each type of crossover, orientation, and genotype separately. The average distance and number of crossovers in each group are indicated below the graph. The simple crossover exchanges were significantly farther from the hotspot center than either the mixed or complex (Mann-Whitney test) while the mixed and complex exchanges were never significantly different from each other. The simple crossover products have exchanges that are, on average, 2-fold closer to the hotspot center than mixed or complex crossover products in *Msh2*^−/−^ mice.

**Table S1.**
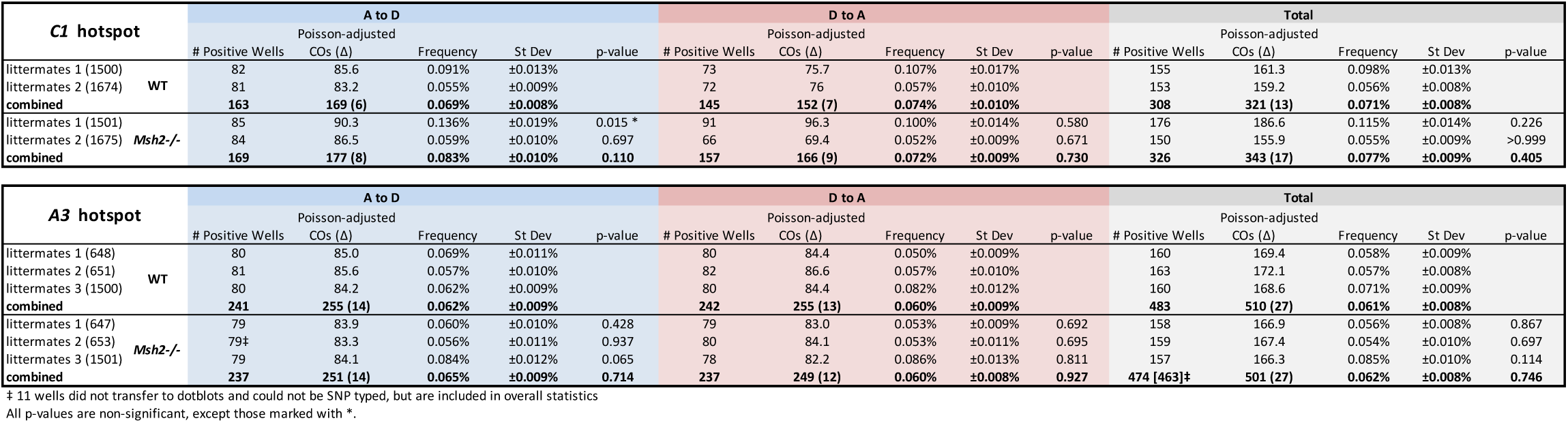
Crossover data and statistics for both hotspots.

**Table S2.** Noncrossover data and statistics for both hotspots.

| C1 hotspot | A chromosome |  |  |  |  | D chromosome |  |  |  |  | Total |  |  |  |  |
| --- | --- | --- | --- | --- | --- | --- | --- | --- | --- | --- | --- | --- | --- | --- | --- |
|  | # Positive Wells (total tracts) | Poisson-adjusted NCOs | Frequency | St Dev | p-value | # Positive Wells (total tracts) | Poisson-adjusted NCOs | Frequency | St Dev | p-value | # Positive Wells (total tracts) | Poisson-adjusted NCOs | Frequency | St Dev | p-value |
| littermates 1 (1500) | 12 | 12.5 | 0.476% | ±0.142% |  | 15 | 15.7 | 0.591% | ±0.159% |  | 27 | 28.2 | 0.534% | ±0.110% |  |
| littermates 2 (1674) | 10 | 10.3 | 0.205% | ±0.070% |  | 13 | 13.5 | 0.268% | ±0.082% |  | 23 | 23.8 | 0.236% | ±0.058% |  |
| combined | 22 (24) | 23 | 0.297% | ±0.067% |  | 28 (28) | 29 | 0.380% | ±0.077% |  | 50 (52) | 52 | 0.338% | ±0.054% |  |
| littermates 1 (1501) | 8 | 8.2 | 0.329% | ±0.119% | 0.505 | 16 | 16.8 | 0.460% | ±0.120% | 0.473 | 24 | 25 | 0.405% | ±0.088% | 0.399 |
| littermates 2 (1675) | 15 | 15.7 | 0.312% | ±0.090% | 0.424 | 21 | 22.4 | 0.445% | ±0.113% | 0.229 | 36 | 38.1 | 0.378% | ±0.080% | 0.117 |
| combined | 23 (23) | 24 | 0.316% | ±0.070% | 0.882 | 37 (37) | 39 | 0.451% | ±0.094% | 0.619 | 60 (60) | 63 | 0.387% | ±0.071% | 0.567 |

| A3 hotspot | A chromosome |  |  |  |  | D chromosome |  |  |  |  | Total |  |  |  |  |
| --- | --- | --- | --- | --- | --- | --- | --- | --- | --- | --- | --- | --- | --- | --- | --- |
|  | # Positive Wells (total tracts) | Poisson-adjusted NCOs | Frequency | St Dev | p-value | # Positive Wells (total tracts) | Poisson-adjusted NCOs | Frequency | St Dev | p-value | # Positive Wells (total tracts) | Poisson-adjusted NCOs | Frequency | St Dev | p-value |
| littermates 1 (648) | 15 | 15.7 | 0.315% | ±0.088% |  | 29 | 31.8 | 0.620% | ±0.135% |  | 44 | 47.5 | 0.466% | ±0.088% |  |
| littermates 2 (651) | 15 | 15.7 | 0.267% | ±0.077% |  | 26 | 28.2 | 0.486% | ±0.105% |  | 41 | 43.9 | 0.373% | ±0.067% |  |
| littermates 3 (1500) | 27 | 29.4 | 0.669% | ±0.144% |  | 24 | 25.9 | 0.550% | ±0.119% |  | 51 | 55.3 | 0.620% | ±0.098% |  |
| combined | 57 (60) | 61 | 0.401% | ±0.069% |  | 79 (85) | 86 | 0.549% | ±0.088% |  | 136 (145) | 147 | 0.475% | ±0.068% |  |
| littermates 1 (647) | 14 | 14.6 | 0.321% | ±0.093% | >0.999 | 48 | 56.5 | 1.122% | ±0.205% | 0.029 * | 62 | 71.1 | 0.714% | ±0.122% | 0.051 |
| littermates 2 (653) | 20 | 21.3 | 0.205% | ±0.053% | 0.482 | 42 | 48.4 | 0.588% | ±0.106% | 0.624 | 62 | 69.7 | 0.368% | ±0.068% | 0.920 |
| littermates 3 (1501) | 15 | 15.7 | 0.352% | ±0.095% | 0.045 * | 34 | 38.0 | 0.830% | ±0.155% | 0.187 | 49 | 53.7 | 0.586% | ±0.094% | 0.841 |
| combined | 49 (51) | 52 | 0.265% | ±0.048% | 0.040 * | 124 (144) | 143 | 0.798% | ±0.114% | 0.029 * | 173 (195) | 195 | 0.509% | ±0.070% | 0.689 |

All p-values are non-significant, except those marked with \*.

**Table S3:**
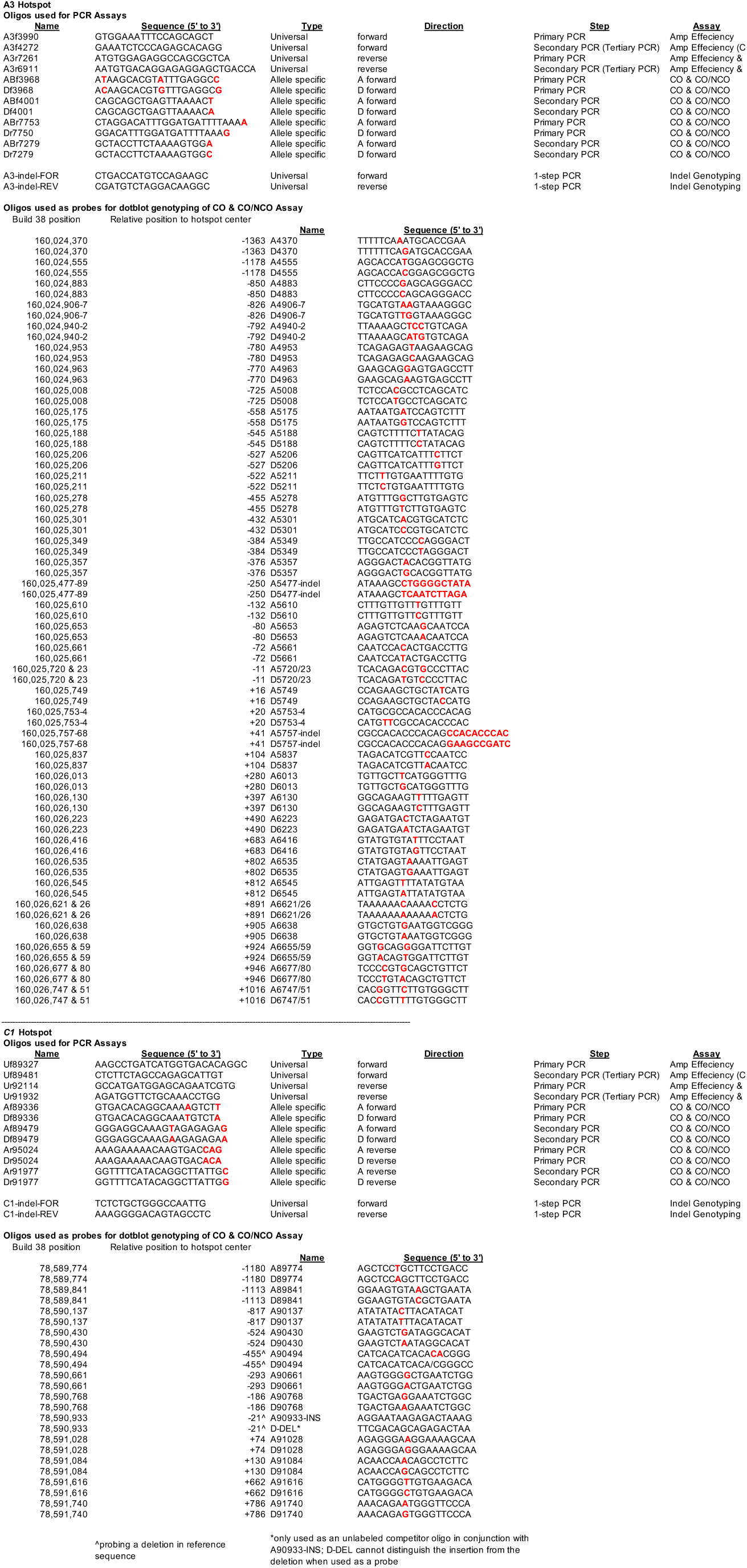
List of allele-specific and universal oligonucleotides used as PCR primers, and oligonucleotides used for Southern blot probes.

## STAR METHODS

### KEY RESOURCES TABLE

STAR METHODS KEY RESOURCES TABLE
| REAGENT or RESOURCE | SOURCE | IDENTIFIER |
| --- | --- | --- |
| **Antibodies** |  |  |
| mouse anti-MLH1 | BD Biosciences | 554073; RRID:AB_395227 |
| rabbit anti-SYCP3 | Abcam | ab15093; RRID:AB_301639 |
| goat anti-mouse A488 | Thermo | A-11001; RRID:AB_2534069 |
| goat anti-rabbit A594 | Thermo | A-11012; RRID:AB_2534079 |
| **Polymerase Chain Reaction Enzymes & Reagents** |  |  |
| Taq DNA polymerase | Thermo | Cat # EP0406 |
| Pfu Turbo DNA polymerase | Agilent | Cat # 600254 |
| Non-acetylated ultra-pure bovine serum albumin (BSA) | Ambion | Cat # AM2616 |
| dNTPs, PCR-grade | Roche | Cat # 3622614001 |
| **Experimental Models: Organisms/Strains** |  |  |
| *Msh2<sup>tm1Rak/+</sup>* heterozygous mice (referred to as *Msh2<sup>+/-</sup>*) | Smits, et. al., 2000. | RRID:MGI:4355514 |
| Inbred A/J mice for backcrossing *Msh2<sup>+/-</sup>* mice | The Jackson Laboratory | Strain # 000646;<br>RRID:IMSR_JAX:000646 |
| Inbred DBA/2J mice for backcrossing *Msh2<sup>+/-</sup>* mice | The Jackson Laboratory | Strain # 000671;<br>RRID:IMSR_JAX:000671 |
| **Oligonucleotides** |  |  |
| All oligos used for PCR assays and Southern dot blotting, See Supplemental Information, Table 3. | Integrated DNA Technologies (IDT), Eton Biosciences |  |
| **Software and Algorithms** |  |  |
| Data analysis and graphical presentation | GraphPad Prism 7 | RRID:SCR_002798 |
| Quantification of dot blots | Image J | RRID:SCR_003070 |
| Calculation of recombination frequencies and other data | Microsoft Excel | RRID:SCR_016137 |

### LEAD CONTACT AND MATERIALS AVAILABILITY

Further information and requests for mouse strains and reagents should be directed to and will be fulfilled by the Lead Contact, Maria Jasin.

### EXPERIMENTAL MODEL AND SUBJECT DETAILS

#### Amplification efficiency and crossover assays from *Msh2*^*–/–*^ mice at the *A3* hotspot (see table below)

The amplification efficiency for each sperm DNA sample was tested by seeding each well with 12 pg DNA, which is on average two copies of A and two copies of D target sequences (4 sperm total). Amplification is performed with each A-S oligo set being tested and an opposing universal oligo, in nested PCR reactions. The amplification adjustment factor (AAF) was determined for each A-S oligo set and DNA sample, in some cases twice, as indicated, to refine the AAF. For the crossover assay— either A to D (rows shaded blue) or D to A (rows shaded red)—the lower value of the AAF from the two A-S oligo sets is used to calculate the amount of input DNA to give 200 amplifiable haploid genomes per well (bold, shaded yellow). In the cases where the amplification efficiency assay was repeated, the amplifiable haploid genomes per well calculated from the seeded DNA per well was adjusted to reflect the new AAF.

The number of wells containing at least one crossover product (and the total number of wells assayed) is used to calculate μ, the average frequency of positive wells; the crossover frequency is calculated by dividing μ by the number of haploid genomes seeded in each reaction. The percent positive wells containing >1 crossover product is estimated based on the Poisson approximation and the value of μ. The percentage of wells containing crossover products with mixed polymorphisms (inferred) is the total percent of positive wells with a mixed crossover product minus the Poisson approximation of the percentage of positive wells containing more than one crossover product.

#### Estimate of crossover products arising from intermediates with heteroduplex DNA

As is seen in the graph below, the inferred percentage of wells containing crossover products with mixed polymorphisms increases somewhat with the amplification efficiency. With an amplification efficiency of 1.0, the percentage of wells containing crossover products with mixed polymorphisms, and hence heteroduplex DNA, is extrapolated to be 46.5%. Complex crossover products are postulated to arise from recombination intermediates with heteroduplex DNA; those without mixed polymorphisms are only 10.6% of total (49 / 463). Thus, we estimate that crossover products arising from intermediates with heteroduplex DNA are maximally 57.1% of total.

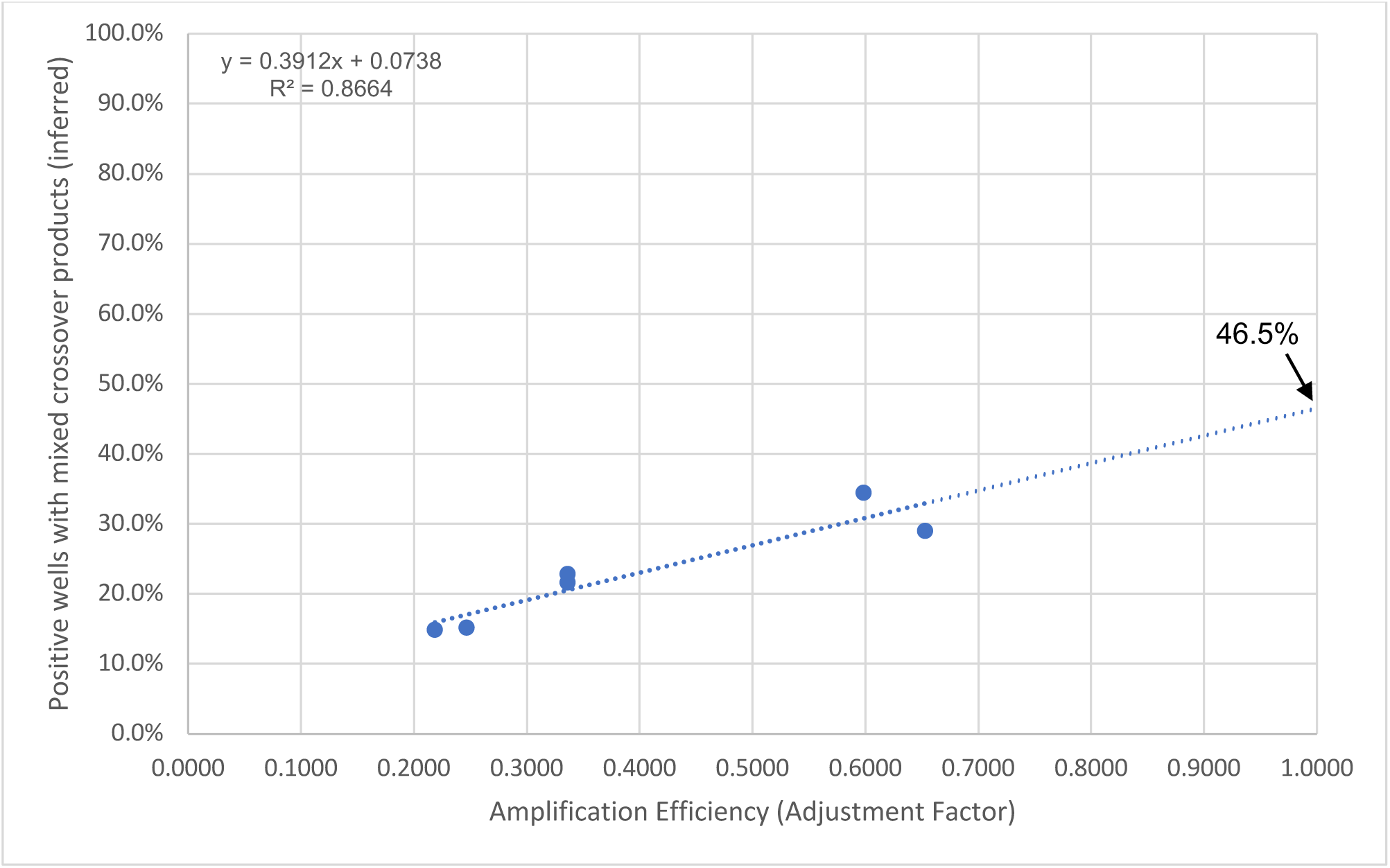

| Amplification Efficiency Assay |  |  |  | Crossover Assay |  |  |  |  |  |  |  |  |  |  |  |
| --- | --- | --- | --- | --- | --- | --- | --- | --- | --- | --- | --- | --- | --- | --- | --- |
| Mouse ID | A-S oligo set<sup>1</sup> | Neg / total wells (# of assays) | AAF<sup>2</sup> | DNA seeded per well (ng)<sup>3</sup> | Haploid genomes /well | Crossover +ve (total wells) | Crossover frequency<sup>4</sup> ($\mu$)<sup>5</sup> | Simple wells | Mixed wells | Complex with mixed wells | Complex without mixed wells | % Simple | % Total mixed<sup>6</sup> | % +ve wells with >1 crossover (Poisson) | % Total mixed<sup>7</sup> (inferred) |
| 647 -/- | Afor | 60/143 (2) | 0.434 | 3.57 | 210.3 | 79 (665) | 0.060% (0.126) | 55 | 19 | 3 | 2 | 69.6 | 27.8 | 6.2 | 21.6 |
| 647 -/- | Drev | 71/144 (2) | 0.354 |  |  |  |  |  |  |  |  |  |  |  |  |
| 647 -/- | Dfor | 57/144 (2) | 0.463 | 3.57 | 194.0 | 79 (808) | 0.053% (0.103) | 49 | 16 | 6 | 8 | 62.0 | 27.8 | 5.1 | 22.8 |
| 647 -/- | Arev | 75/144 (2) | 0.336 |  |  |  |  |  |  |  |  |  |  |  |  |
| 653 -/- | Afor | 34/95 (1) | 0.514 | 4.86 | 200.0 | 68<sup>8</sup> (747) | 0.056% (0.112) | 49 | 12 | 2 | 5 | 72.1 | 20.6 | 5.5 | 15.1 |
| 653 -/- | Drev | 58/95 (1) | 0.247 |  |  |  |  |  |  |  |  |  |  |  |  |
| 653 -/- | Dfor | 60/192 (2) | 0.368 | 5.49 | 200.0 | 80 (802) | 0.053% (0.105) | 41 | 9 | 7 | 23 | 51.3 | 20.0 | 5.2 | 14.8 |
| 653 -/- | Arev | 62/96 (1) | 0.219 |  |  |  |  |  |  |  |  |  |  |  |  |
| 1501 -/- | Afor | 44/288 (3) | 0.963 | 1.42 | 157.2 | 79 (640) | 0.084% (0.132) | 50 | 27 | 1 | 1 | 63.3 | 35.4 | 6.4 | 29.0 |
| 1501 -/- | Drev | 51/192 (2) | 0.653 |  |  |  |  |  |  |  |  |  |  |  |  |
| 1501 -/- | Dfor | 56/288 (3) | 0.673 | 1.42 | 126.6 | 78 (756) | 0.086% (0.109) | 37 | 20 | 11 | 10 | 47.4 | 39.7 | 5.3 | 34.4 |
| 1501 -/- | Arev | 66/192 (2) | 0.599 |  |  |  |  |  |  |  |  |  |  |  |  |

<sup>1</sup>See Table S3 for allele-specific (A-S) oligo sequences.
<sup>2</sup>AAF, amplification adjustment factor (amplification efficiency); $[-\ln(\text{negative wells} / \text{total wells}) / 2]$ .
<sup>3</sup>Based on amplification efficiency.
<sup>4</sup>Crossover frequency in sperm [ $\mu$ / haploid genomes per well].
<sup>5</sup>Mean frequency of positive wells $[-\ln(\text{negative wells} / \text{total wells})]$ .
<sup>6</sup>Mixed wells + Complex with mixed wells / total crossover positive wells.
<sup>7</sup>% Total mixed minus % +ve wells inferred to have >1 crossover, based on Poisson approximation.
<sup>8</sup>Of 79 positive wells, only 68 were genotyped.

### METHOD DETAILS

#### Mouse care

The care and use of mice in this study were performed in accordance with a protocol approved by the Institutional Animal Care and Use Committee (IACUC) at Memorial Sloan Kettering Cancer Center (MSKCC). Mice were housed under Federal regulations and policies governed by the Animal Welfare Act (AWA) and the Health Research Extension Act of 1985 in the Research Animal Resource Center (RARC) at MSKCC, and was overseen by IACUC.

#### Mice husbandry and genotyping

Wild-type inbred mice were obtained from Jackson Laboratory (A/J stock #000646, DBA/2J stock #000671). *Msh2*^*tm1Rak/+*^ mice were a gift from W. Edelmann (Smits et al., 2000) and genotyped with a three primer system as described previously (Kovalenko et al., 2012). These mice were originally on a C57BL/6 and 129S mixed background. The *A3* locus was genotyped by restriction digest of two amplicons, Left and Center, which can distinguish the 4 inbred strains: Left amplicon: Chromosome 1: 160,023,339 −160,025,391 (Relative position: −2394 bp to −342 bp)

A3f3339: 5’ TGTGTCAGGTGAAATAAGGCA 3’

A3r5391: 5’ TCAGTCAGTTGTCAGAGACC 3’

Center amplicon: Chromosome 1: 160,025,337-160,026,193 (Relative position: −396 bp to +479 bp)

A3f5337: 5’ TGTTGCCATCCCCAGGGACT 3’

A3r6193: 5’ GATTTGGCCAACATTGTGGG 3’

The Left amplicon is digested with StuI and SacI, which produces these approximate fragment sizes:

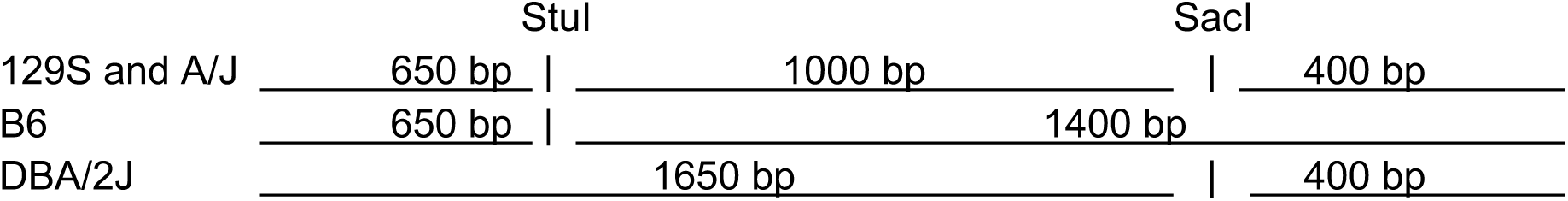

The Center amplicon is digested with DraI (optional) and HhaI, which produces these approximate fragment sizes:

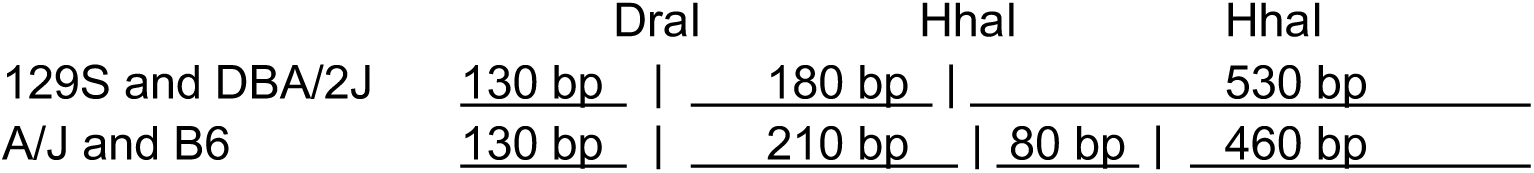

For *A3* analysis, littermates #1 (647s & 648) and #2 (651 & 653) were from different litters of the same parents: the DBA/2J sire was backcrossed twice (A3 was genotyped at first and second backcross, and was homozygous D/D), the A/J dam was backcrossed three times (A3 was verified A/A by the second backcross). The genotype at *A3* was verified in tail DNA from the experimental F1 hybrids by sequencing each allele separately. The third set of experimental littermates (1500 & 1501) was obtained from inbred dam (D/D) and sire (A/A) which were backcrossed more than ten times, and therefore congenic.

The third set of experimental hybrids was also used for the *C1* hotspot analysis (1500 & 1501). Likewise, the second set of experimental hybrids used for *C1* analysis were obtained from inbred dam (D/D) and sire (A/A) that were backcrossed more than ten times (1674 & 1675).

#### Isolation of sperm DNA

Sperm DNA from mice 647, 648, 651 and 653 was performed as described previously (Cole and Jasin, 2011). Sperm DNA from mice 1500, 1501, 1674 & 1675 we isolated by a modified, gentler method which produces high quality and high yield sperm DNA with virtually no somatic cell contamination, without enzymatic tissue digestion or differential SDS-lysis of somatic cells. Both cauda and caput epididymides were dissected from >2 month old mice, with all excess fat and tissue trimmed away. Epididymides were cut in quarter chunks and placed in the cap of 5mL tube with cell-strainer cap (BD # 352235) filled past the mesh with PSB, such that the tissue is placed on top of the filter cap but in a continuous buffer with the tube. The sperm swim down out of the tissue, through the filter cap and into the tube by gravity. Let sit for 5-10 minutes, and carefully lift the cap out of the PBS to break the fluid surface tension, which will release the sperm into the tube. Do this every 5 minutes until sperm are no longer released upon breaking the surface tension. Remove the cap (and tissue) and seal the tube with parafilm. Pellet the sperm in a swing-bucket rotor for 2 minutes at 4,000 x g. Aspirate the supernatant down to 1 mL, resuspend the sperm by gentle vortex pulses and transfer to 2 mL screw-cap tube. Remove 5 μL and place on slide to access sperm quality and lack of somatic cell contamination. Pellet in a swing-bucket rotor for 5 minutes at 4,000 x g, and completely aspirate supernatant. Add 600 μL of lysis buffer (200 mM NaCl, 100 mM Tris-Cl pH 7.5, 5 mM EDTA, 0.5% SDS, 1.5 M 2-mercaptoethanol, 0.5 mg/mL Proteinase K). Briefly and gently vortex with quick pulses to resuspend. Incubate at 55°C for 3-4 hours. Extract DNA with 600 μL phenol:choloroform, remove aqueous phase and repeat extraction again with 600 μL phenol:chloroform. Place the aqueous phase in a new tube and precipitate DNA with 1.2 mL 100% ethanol. Spin and wash pellet with 0.75 mL 70% ethanol. Spin and resuspend the pellet in 100 μL 5 mM Tris-Cl, pH 7.5. Incubate at 55°C for 1 hour, then overnight at 4°C. Quantify DNA by gel electrophoresis and OD_260_. Store DNA in 10 μL aliquots and freeze at −20 °C, or −80°C for long-term storage.

#### Crossover and Noncrossover assays

Experiments were performed as described previously (Cole and Jasin, 2011; Cole et al., 2010). Preliminary experiments are carried out to assess the quality of the template and each allele-specific oligo, by seeding reactions with two copies of each parental haplotype (12 pg hybrid sperm DNA), with an allele-specific oligo and a “universal” oligo which hybridizes with both haplotypes. For a complete list of all PCR oligonucleotides and Southern probe oligonucleotides used, see Table S3. A Poisson distribution function is applied to determine the number of amplifiable templates seeded in each well, and this adjustment factor is applied to all subsequent PCRs using the same allele-specific oligo and sperm DNA sample (see Supplemental Discussion).

All PCR reactions use 10X “Jeffreys” buffer: 450 mM Tris, pH8.8, 110 mM (NH_4_)_2_SO_4_, 45 mM MgCl_2_, 67 mM 2-mercaptoethanol, 44 μM EDTA, 10 mM each dNTP (Roche, Cat # 3622614001), 1.13 mg/mL ultra-pure (non-acetylated) BSA (Ambion Cat # AM2616), 12.5 mM Tris base (not pH adjusted). Final reaction conditions are as follows: 1X Jeffreys buffer, 0.2 μM each primer, 0.03 U/μL Taq (Thermo Cat # EP0406), 0.006 U/μL Pfu Turbo (Agilent Cat # 600254), in a 10 μL volume. See (Cole and Jasin, 2011) for experimental details, including nested cycling conditions, S1-nuclease digestion, and all other aspects of allele-specific PCR assays and polymorphism genotyping.

To combine data from multiple mice, the Poisson-adjusted crossovers and noncrossovers, and total DNA assayed in each interval from each dataset were summed, with subsequent frequencies and cumulative fractions calculated from these sums after this initial Poisson correction.

Tract lengths are always listed as minimum lengths from the first to the last involved polymorphisms.

#### Clonal analysis of DNA from positive crossover wells

DNA from crossover PCR reactions containing *Msh2*^*–/–*^ DNA which were found to contain mixed or complex tracts by Southern dotblotting was TOPO-cloned. Thirty-two individual clones were patch-streaked only agar plates with control clones containing the same amplicon of non-recombinant A or D sequence. Bacteria was replica plated onto several Nylon membranes, cells were lysed by laying the membrane on filter paper soaked in 0.5M NaCl, 0.5M NaOH for 2 minutes and neutralized by laying on filter paper soaked in 1.5 M NaCl, 0.5 M Tris-Cl pH 7.5. The DNA was fixed by drying at room temperature and crosslinking with a Stratalinker. Southern hybridization was performed as described for CO and NCO assays. Note: Some clones were apparently empty vector, while other clones did not transfer, lyse, or became mixed with neighboring patches. Therefore, the total number of clones analyzed is quite variable.

#### Spermatocyte chromosome spreads and Immunofluorescence analysis

Testes were collected from mice 1500 (WT) & 1501 (*Msh2*^*–/–*^) and spermatocytes were prepared for surface spreading as previously described (Barchi et al., 2008). SYCP3 (rabbit anti-SYCP3; ab15093; RRID:AB_301639; 1:500) and MLH1 (mouse anti-MLH1; BD Biosciences 554073; RRID:AB_395227; 1:25) were detected by diluting the primary antibodies in dilution buffer (0.2% BSA, 0.2% fish gelatin, 0.05% Triton X-100, 1 x PBS) and incubating overnight at 4 °C. Slides were subsequently washed in dilution buffer three times and incubated with the following secondary antibodies at 1:200 dilution for 1 h at 37 °C: A594 goat anti-rabbit (Thermo A-11012; RRID:AB_2534079) and A488 goat anti-mouse (Thermo A-11001; RRID:AB_2534069). Cover slips were mounted with ProLong Gold antifade reagent with DAPI (Invitrogen P36935).

### QUANTIFICATION AND STATISTICAL ANALYSIS

#### Scoring Southern dot blots for crossover and noncrossover genotyping

Each dot blot includes a control dilution series of the haplotype being probed: 100%, 1:10, 1:30, 1:100, 1:300, and 1:1000 of the control haplotype DNA is diluted into DNA of the opposite haplotype. Blots are imaged with a phosphoimager (GE Typhoon 7000), and the intensity of each dot is quantified in Image J (https://imagej.nih.gov/ij/). A standard curve is created, and the intensity of each experimental dot is compared to this curve, with only signals surpassing the 1:30 dilution being considered positive.

All statistical tests are performed using GraphPad Prism software (https://www.graphpad.com), and the name of the test is indicated in the text or figure legend.

